# SPTnet: a deep learning framework for end-to-end single-particle tracking and motion dynamics analysis

**DOI:** 10.1101/2025.02.04.636521

**Authors:** Cheng Bi, Kevin L. Scrudders, Yue Zheng, Maryam Mahmoodi, Shalini T. Low-Nam, Fang Huang

## Abstract

Single-particle tracking (SPT) provides high-resolution spatial-temporal information on biomolecule dynamics. However, localization inaccuracies, limited track lengths, heterogeneous fluorescence backgrounds, and potential molecular motion blur pose significant challenges that hinder the accurate extraction of movement trajectories and their underlying motion behavior. The conventional SPT pipeline struggles to comprehensively address detection, localization, linkage, and motion parameter inference simultaneously, resulting in information loss during sequential processing. To overcome these challenges, we propose SPTnet, an end-to-end deep learning framework that leverages a Transformer-based architecture to optimize trajectory and motion parameter estimations in parallel through a global loss. SPTnet bypasses traditional SPT processes, directly inferring molecular trajectories and motion parameters from fluorescence microscopy videos with a precision approaching the statistical information limit. Our results demonstrate that SPTnet outperforms conventional methods under commonly encountered but challenging conditions such as short trajectories, low signal-to-noise ratio (SNR), heterogeneous backgrounds, motion blur, and especially when molecules exhibit non-Brownian behaviors.

## INTRODUCTION

Single-particle tracking (SPT) is a powerful tool for studying subcellular dynamics in living systems. By tagging molecules of interest with fluorescent probes, it enables the observation of the dynamic motion behavior of a single molecule in live cells with high spatial and temporal precision^1^. In contrast to ensemble measurements such as fluorescence recovery after photobleaching (FRAP)^2^ and fluorescence correlation spectroscopy (FCS)^3^, SPT provides unique insights into the motion behavior of individual molecules, revealing subcellular and molecular heterogeneities and interactions that are often lost in ensemble averages.

For a freely diffusing molecule in anisotropic surroundings, Brownian motion is a suitable mathematical model^4^. However, in the complex intracellular environment, non-Brownian motion behaviors are often encountered, as observed in a diverse set of biomolecules and biological model organisms, such as, mRNA in *E. coli*^5^, telomeres in the mammalian cells^6^ and melanosomes in *Xenopus laevis*^7^. Quantitative understanding of these complex anomalous motion behaviors requires mathematical models^8^ that describe the rules governing these biomolecular movements such as Continuous-time random walk (CTRW), Lévy walk (LW), and fractional Brownian motion (fBm). Specifically, fBm enables the distinction of different motion types through its parameterized Hurst exponent, characterizing subdiffusion, Brownain motion, and superdiffusion. This model provides a generalized framework for modeling both persistent and anti-persistent movement, which can arise from the heterogeneous cellular environment or biological activities, such as spatial protein gradients in embryo development^9^ and intracellular cargo transport^10, 11^

Besides these mathematical models, the precise retrieval of single-molecule positions, their trajectories over time, and associated motion parameters from experimental data are essential for gaining meaningful insights into biomolecular movements. However, this process is often challenging and prone to errors at each step of the conventional SPT analysis pipeline. First, particle detection and localization rely heavily on empirical parameter choices and often perform inconsistently under low signal-to-noise conditions, such as in live-cell experiments. In addition, heterogeneous backgrounds induced by autofluorescence and out-of-focus emitters often cause artifacts, even during the initial localization step^12^.

Second, conventional trajectory linking algorithms mainly rely on consecutive time-lapse frames, whereas long-range correlated motion behaviors inherent to some biomolecules are often challenging to consider^13, 14^. While Bayesian-based approaches are capable of considering global information to optimize the joint posterior distribution of tracks, these methods often require a robust prior, such as the probe’s photophysical properties and the transition probability of motion behaviors, in addition to a reliance on empirical parameters such as the maximum searching radius. Their practical applications are further hindered by the significant computational complexity – rapidly escalating computation time with the number of molecules and camera frames.

Even with perfectly reconstructed trajectories, extracting motion dynamic information may be challenging. The widely used mean square displacement (MSD) analysis requires long trajectories for robust estimates due to its asymptotic nature. Moreover, it is difficult to satisfy the criteria and prerequisites to properly fit the MSD plot for most experimental situations^15^.

Recent studies have demonstrated the capability of convolutional neural networks (CNNs) and recurrent neural networks (RNNs) for estimating the Hurst exponent and inferring the underlying motions from short trajectories^16,17,18^. The performance of these methods, however, relies on accurately constructed particle trajectories, which are assumed to be free of errors in detection, localization, and linkage, a condition that is challenging to meet practically. To date, no existing end-to-end, parameter-free method enables the tracking and extraction of dynamic information about molecules directly from camera images with high accuracy.

Here, we propose SPTnet, a transformer-based deep learning framework for single particle tracking and motion dynamics analysis. SPTnet directly processes camera videos of moving particles and outputs their estimated trajectories, generalized diffusion coefficients and Hurst exponents with precision approaching the theoretical limit. SPTnet bypasses the traditional processes such as particle detection, localization, trajectory linkage, and regression, eliminating information loss and potential biases introduced by conventional algorithms. Leveraging a *de novo* Transformer-based architecture, proven effective in revolutionary large language models like ChatGPT, SPTnet processes all frames in parallel, providing access to global spatial-temporal information. We show that this holistic view of the SPT pipeline mitigates overall information loss with a precision approaching the theoretical information limit. We demonstrate that SPTnet outperforms conventional methods under challenging conditions including short trajectories, low SNR, heterogeneous background, and motion blur. We applied SPTnet to experimental datasets collected from three distinct imaging systems. First, we validated the method by tracking fluorescent beads in glycerol-water mixtures, achieving results consistent with theoretical calculations. We then analyzed low-SNR data on supported lipid bilayers (SLBs) and successfully identified both immobile and Brownian motion populations. Furthermore, we applied SPTnet to single-particle tracking photoactivated localization microscopy (sptPALM) and unraveled the distinct motion behaviors of Reticulon 4 (Rtn4) and Sec61β along endoplasmic reticulum (ER) tubules in live COS-7 cells.

## RESULTS

### Design of SPTnet

SPTnet is designed to analyze the motions over time of single fluorescent molecules within sequences of light microscopy images. Rather than following traditional sequential processing pipelines, SPTnet simultaneously addresses localization, tracking, and motion parameter inference as a set prediction problem. Inspired by the success of the Transformer^19^ in image classification and segmentation tasks^20–22^, the architecture of the SPTnet comprises of following components: a customized 3D-ResNet backbone, two-stream decoder-encoder transformers, feature fusion module, and a 4-headed multilayer perceptron (MLP) (**Supplementary Note 1**). This approach enables end-to-end propagation, allowing the model to directly predict each particle’s trajectory, along with its corresponding motion parameters from a video (**Fig. 1a**).

**Figure 1.**
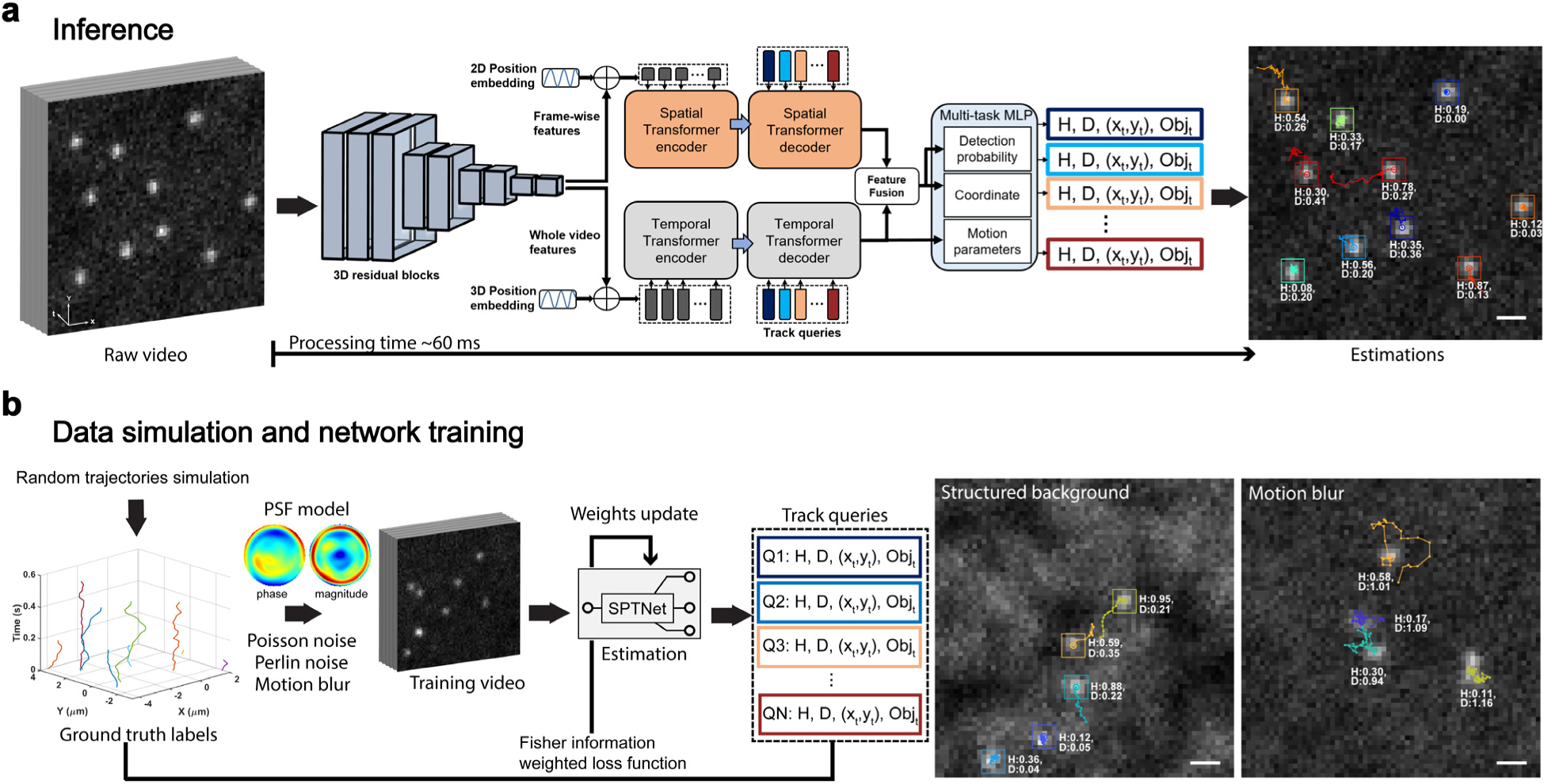
End-to-end single particle tracking through video using SPTnet. **a,** Architecture and inference pipeline of SPTnet. Sequences of images can be directly analyzed by the transformer-based neural network, which simultaneously outputs trajectories, Hurst exponent (H), and generalized diffusion coefficient (D) for each molecule within the video. **b,** Training workflow of SPTnet with example training frames, showing structured backgrounds and motion blur, overlaid with ground truth trajectories and corresponding motion parameters. Training videos for SPTnet are simulated using a pupil function with various noise sources. For details, see Supplementary Note 4. Scale bar, 1 µm.

Specifically, the backbone extracts feature maps from sequences of single-molecule emission patterns in recorded camera frames using 3D convolutional layers with residual connections. The 3D convolutional kernels capture spatial-temporal features, making SPTnet capable of distinguishing moving particles that are spatially overlapped in certain frames but separable in the temporal domain (**Supplementary Fig. 2 and Supplementary Video 1**). Then the two-stream transformers module processes the extracted video features by assigning weights based on their significance and relevance to each track through the attention mechanism. Within this module, one transformer (Spatial-T) focuses on predicting the spatial position of the particles in each frame, while the other (Temporal-T) processes all the frames together to capture motion dynamics information.

A feature-fusion module then combines information from both streams, producing integrated position and motion features. This integration allows Temporal-T to leverage global information by assisting Spatial-T to recognize and link emission patterns that are ambiguous solely based on the decoding using frame-wise information. The fused features, along with direct outputs from Temporal-T are then sent to the final multi-task MLP, which simultaneously generates estimations for detection probability, particle coordinates, Hurst exponent, and generalized diffusion coefficient (**see Supplementary Fig. 3 and Supplementary Table 1 for detailed architecture of SPTnet**). This design is not constrained by localizations within a fixed grid or predefined anchor boxes in each frame, offering flexibility for handling videos with varying numbers of molecules, different start times, durations, and diverse motion types.

### Loss function and track specific information weighting

We trained SPTnet to estimate trajectories and motion parameters simultaneously by constructing a global loss function. This loss function is a linear combination of four components: (1) mean absolute error of the Hurst exponent, (2) generalized diffusion coefficient, (3) binary cross-entropy loss for the probability that a molecule is present in each frame, and (4) Euclidean distance loss for the coordinates. By incorporating information about motion dynamics into the loss function, the network is trained to reconstruct trajectories based on both estimated molecule positions and their motion behaviors. This approach reduces ambiguities that often cause false linkages among spatially overlapped molecules. To correctly associate each ground truth labeled molecule with a track query for loss calculation, we used the Hungarian algorithm^23^ for bipartite assignment. For unmatched track queries, the detection loss of background is calculated and added to the integrated loss of matched molecules for backpropagation (**Supplementary Note 5**). Therefore, all the molecules in a video can be estimated together, with each track query forced to follow one molecule, thereby eliminating redundant detection of the same molecule.

Optimization of this multi-task loss function among various video data can be challenging. A single video may contain multiple molecular tracks that vary in length and motion type, each of which differs significantly in inference difficulty. For example, analyzing a short track with fast Brownian motion diffusion is considerably more challenging than analyzing a long track with slow and superdiffusive behavior (**Supplementary Fig. 4**). This variation can reduce the convergence speed and potentially trap the model in suboptimal local minima. To facilitate the training and convergence of SPTnet, we normalize the Hurst exponent and diffusion coefficient losses based on their statistical precision limits, quantified through Fisher information (**Supplementary Note 3**). For instance, molecules undergoing Brownian motion that appear in only a few frames are expected to have low Fisher information content, leading to lower weights in their loss functions and a greater error allowance that matches their statistical precision limits.

### Training data generation

We trained SPTnet with >200,000 simulated videos containing emission patterns from single molecules following stochastic traces of fractional Brownian motion corrupted by photon detection noise, and both uniform and heterogeneous backgrounds. To eliminate discrepancies between simulated and experimental data, single molecule emission patterns were generated based on the pupil function - representing the electric field distribution in the back focal plane of the objective in the microscope system (**Supplementary Note 4.2**). Consequently, PSFs from various imaging systems, including those with system-induced wavefront distortions, can be accurately modeled using the phase-retrieved pupil function derived from fluorescent beads or *in situ* PSF modeling^24, 25^ (**Supplementary Fig. 5**).

Single particle tracking data are often captured in live cells, and in some cases tissues, where intra-and-extracellular structures are distributed across three dimensions both in and out of focus. Such complexities give rise to heterogeneous backgrounds generated by out-of-focus fluorescent labels and autofluorescence from cells and tissues, leading to incorrect estimation of trajectories and motion behaviors in conventional SPT analysis. To this end, SPTnet incorporates the heterogeneous background into its training videos by modeling it with Perlin noise^26^ (**Supplementary Note 4.3**). Since Perlin noise, with its different spatial frequency components, can adapt to various textures, and the smooth interpolation used in the algorithm makes it closely resemble the structured background observed in the experimental data^12, 27^.

Moreover, fast-diffusing molecules will introduce motion-blurred PSFs with finite camera exposure times, which must be accounted for accurately when estimating motion parameters^28^. We trained SPTnet to decode motion-blurred PSFs through incorporating intermediate frames with tenfold shorter exposure times (**Supplementary Note 4.4**). By integrating these frames to match standard exposure durations, we generated realistic training videos that enabled convolutional kernels to learn and extract features from the blurry PSFs of rapidly diffusing molecules.

### SPTnet predicts motion parameters with high precision under various conditions

We first evaluated SPTnet’s performance in estimating motion parameters using 30-frame simulated datasets and compared it to conventional estimators for characterizing diffusion dynamics, including the wavelet version of second order derivative (wDSOD) method^29^, rescaled range (R/S) analysis^30^, and MSD analysis (**Supplementary Note 6**). To provide an optimal performance baseline, we calculated the theoretical precision limits, the Cramér-Rao lower bound (CRLB), accounting for the stochastic nature of particle movement, trajectory length, and the limited localization precision due to Poisson noise (**Supplementary Note 3 and Supplementary Fig. 4**).

We found that SPTnet outperforms conventional algorithms across different Hurst exponents with a fixed diffusion coefficient, and its estimation precision closely approaches the information limit predicted by the CRLB for the Hurst exponent (**Fig. 2a, left**). Compared to SPTnet’s achieved estimation error (12.5%±7.4%, mean±s.d., higher than the theoretical precision limit, 10 Hurst conditions each containing 1000 videos), the performance of wDSOD, exhibited an error of 338.2%±86.7% (Fig. 2a, left) higher than the CRLB predicted value, which is ∼40 times worse than SPTnet. This is due to wDSOD’s slow rate of convergence and its sensitivity to localization imprecision (**Supplementary Fig. 1**). The R/S method showed an estimation error of 142.2%±68.6% (**Fig. 2a, left**) higher than the theoretical precision limit, and this error increased as the estimated Hurst exponent neared 1 (superdiffusion), consistent with previous reports^31–33^. Finally, compared to the MSD analysis, which showed an estimation error of 35.6% ± 27.4% (**Fig. 2a, left**) above the theoretical precision limit, SPTnet performed ∼2.8-fold better. Notably, SPTnet showed significantly improved performance for highly anomalous motion types, e.g. H=0.1 and H=0.9, with 11-fold and 3.6-fold better than MSD analysis, respectively. When estimating motion behaviors approaching Brownian motion, SPTnet showed a slight improvement over MSD analysis, with both methods nearing the theoretical precision limit. We also evaluated SPTnet’s performance in estimating generalized diffusion coefficients with a fixed Hurst exponent (H=0.5) under Brownian motion conditions. The estimation error of SPTnet is 21.5% ± 8.2% (mean ± s.d.) above the theoretical precision limit, which is 2.6 times better than that of the MSD analysis (55.9% ± 5.1%) (**Fig. 2a, right**).

**Figure 2.**
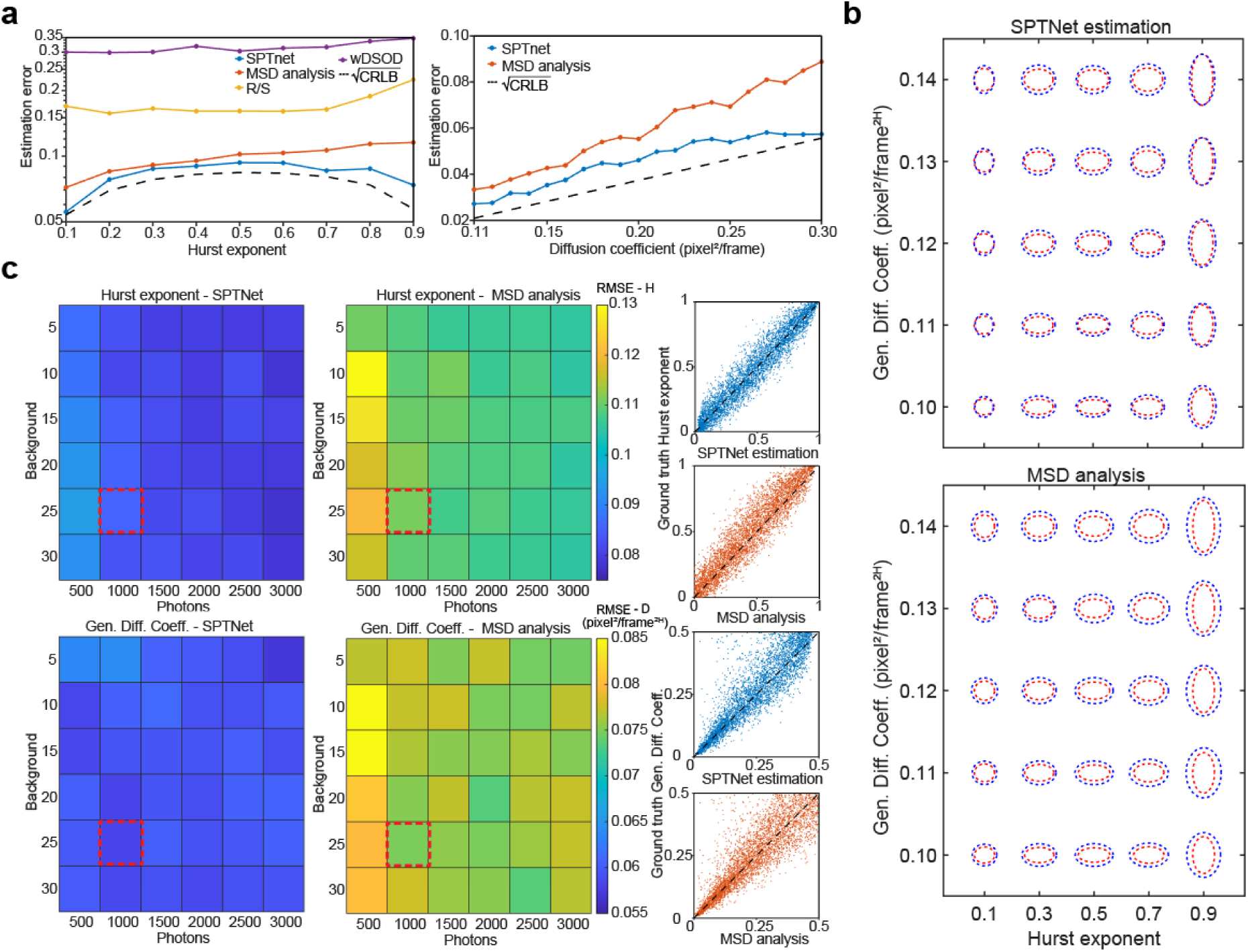
Characterization of the performance of SPTnet on Hurst exponent and generalized diffusion coefficient estimation. **a,** Root mean square error (RMSE) of SPTnet estimation on Hurst exponent compared with three different estimators and theoretical estimation precision limit predicted by the CRLB. For this comparison, 1000 videos were simulated per Hurst exponent. Each video consists of 30 frames, with exactly one molecule per frame. The PSF of the molecule is aberration-free, with constant photon counts of 1,000, background intensity of 25, and generalized diffusion coefficient D=0.25 pixel^2^/frame^2H^ (simulation unit, pixel size: 157 nm, H: Hurst exponent), and the video is corrupted by Poisson noise. The panel on the right shows a similar comparison for diffusion coefficient estimation, given H=0.5 (Brownian motion) for all cases. **b,** Comparison of estimation precision for different combinations of Hurst exponent and generalized diffusion coefficient between SPTnet, MSD analysis, and theoretical precision limit across different motion parameters. The horizontal semi-axis of the red dotted ellipse is the normalized square root of the CRLB of Hurst exponent (normalized to the mean for all 25 plotted cases), and the vertical semi-axis is the normalized square root of the CRLB of the generalized diffusion coefficient. Semi-axes of the Blue dotted ellipses are based on the standard deviation of the estimations from SPTnet (top) and fitting of the MSD plot (bottom). **c,** Comparison of the estimation RMSE for both Hurst exponent and generalized diffusion coefficient across different SNRs. Estimations distribution versus ground truth for the red dotted box SNR is shown on the right (dotted black line indicates perfect estimations). Each SNR condition was tested using 5000 simulated videos, with each containing one moving molecule following the uniformly distributed Hurst exponent and generalized diffusion coefficient. (for all the comparisons, videos with molecules moving outside the field of view were removed from the calculations).

Next, we estimated the error across a diverse combination of Hurst exponents and generalized diffusion coefficients, as different motion types yield varying theoretical precision limits, which can affect the performance of estimators differently. We compared the estimation precision of SPTnet and MSD analysis with the theoretical precision limit. On average, SPTnet estimations are 11.5% and 6.4% larger than theoretical precision limits for the Hurst exponent and generalized diffusion coefficient, respectively. In contrast, the precision of estimates by MSD analysis was worse, 31.2% and 11.4% above the CRLB predicted values (**Fig. 2b**). This high estimation precision allows SPTnet to capture motion behavior switching with high sensitivity (**Supplementary Video 4**).

SPTnet directly infers motion parameters from its video input, making the signal-to-noise ratio (SNR) of the image sequences critical for the network’s performance and its robustness against varying imaging conditions. Therefore, we investigated 36 SNR conditions, each containing 5000 videos. SPTnet consistently had 24.7%±2.4% and 21.9%±7.8% (mean±s.d.) lower errors in estimating Hurst estimation and generalized diffusion coefficient than MSD analysis (**Fig. 2c**). The estimation distribution versus ground truth for photon counts of 1000 and background intensity of 25 (red dotted box) is shown on the right of Figure 2C. In terms of the Hurst exponent, SPTnet estimation has minimal bias with the distribution centered around the black dotted line (indicating perfect estimations), and the width follows the general trend of the predicted precision limit, where information content is at its maximum in highly anomalous motion cases and reaches its minimum in the Brownian motion case. In contrast, the distribution of estimations from MSD analysis is broader and biased toward smaller Hurst exponent values. For the diffusion coefficient, SPTnet estimations show a narrower distribution compared to MSD analysis. Additionally, when the estimated generalized diffusion coefficient approaches the edge of the training range at D=0.5, SPTnet tends to underestimate the value. This bias can be eliminated by extending the range of generalized diffusion coefficient in the training data, though this adjustment may create imbalanced track lengths across motion types and add complexity to the training process, as faster-moving molecules are more likely to exit the field of view within a shorter duration.

### SPTnet enables accurate SPT analysis of videos with highly heterogeneous backgrounds and blurred motions

In diverse and intricate cellular environments, accurate SPT analysis is often compromised by heterogeneous backgrounds and motion blur, which can obscure particle trajectories and lead to inaccurate estimates of motion parameters. Instead of focusing on the often hard-to-interpret emission patterns in single frames (**Fig. 3a and Fig. 3d**), SPTnet leverages global spatial-temporal information extracted from the entire video to distinguish and reconstruct tracks from obscured emitters.

**Figure 3.**
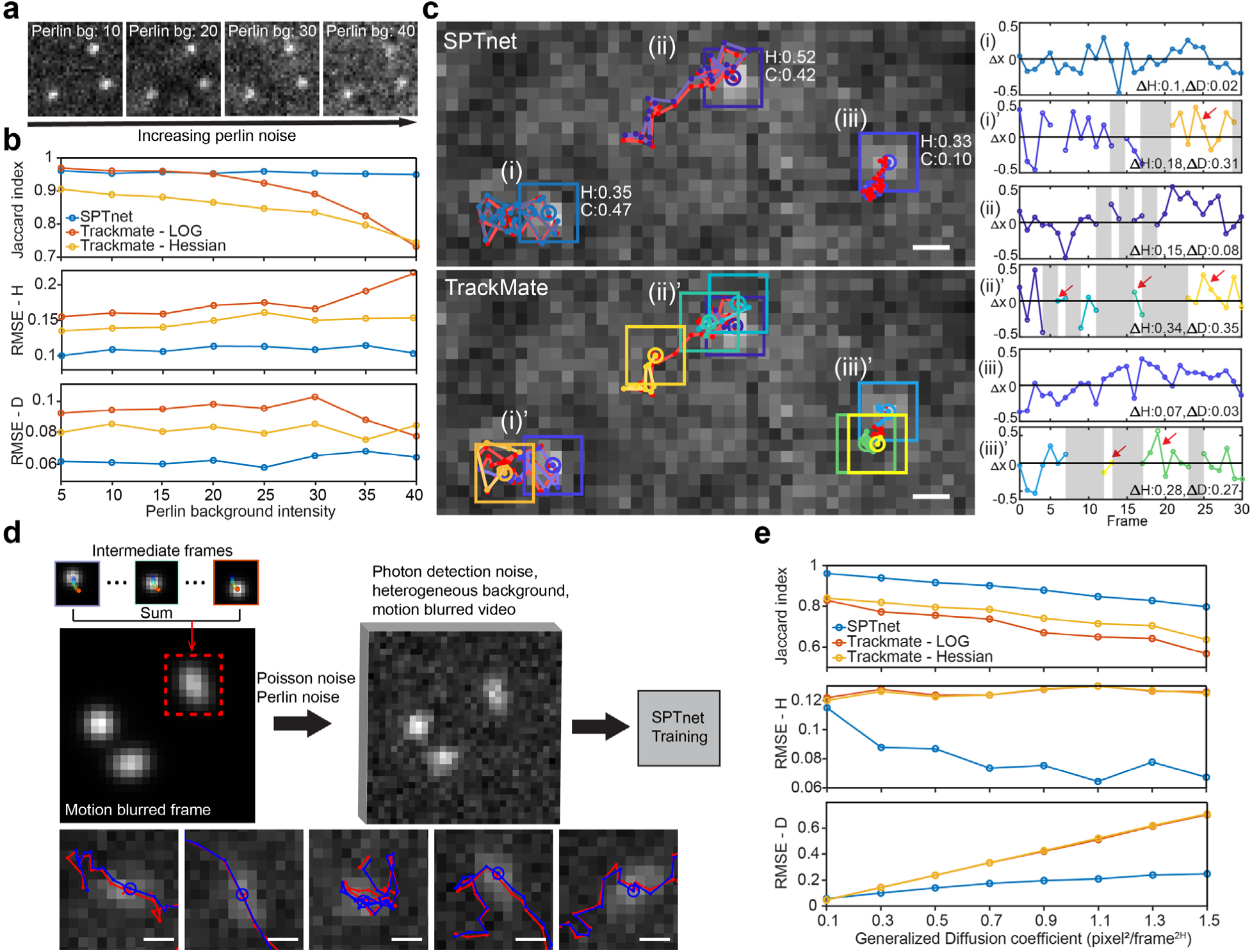
SPTnet enables simultaneous tracking and motion parameter estimation under structured background and motion blur conditions. **a,** Simulated camera frames with increasing levels of Perlin noise, each containing three emitters. **b,** A comparison between SPTnet and TrackMate (using built-in “LoG” and “Hessian” detectors with the LAP tracker, followed by MSD analysis) under different Perlin noise intensity levels. The detailed settings can be found in Supplementary Table 4. We evaluate detection accuracy and estimation error of Hurst exponent (H) and generalized diffusion coefficient (D) by calculating the Jaccard index and RMSE, respectively. Each case was evaluated using 1,000 videos, with each video containing up to five emitters. **c,** Tracking results of Perlin noise intensity at 40, showing ground truth positions in each frame marked with a red line. Different estimated tracks are displayed in various colors. The x-coordinate deviations from the ground truth for each emitter are shown on the right. Shaded areas represent missing detections, red arrows indicate misinterpretations leading to separate tracks, and the absolute estimation error of H and D are shown on the bottom right of each panel. See Supplementary Fig. 7 and Supplementary Video 2 for additional comparisons. **d,** A diagram illustrates the incorporation of motion blur into the training data, along with examples of inference results from frames containing highly blurred PSFs. The ground truth is marked in red, and SPTnet estimations are shown in blue. **e,** Similar comparison to panel b, under conditions of increasing the D (increasing motion blurred effect). Scale bar, 500 nm

In comparison to the rapidly deteriorating performances using the state-of-the-art SPT toolbox, TrackMate^34^, SPTnet maintains its performance in the presence of an extended range of heterogeneous background levels. We compared two built-in detectors in TrackMate: the Laplacian of Gaussian (LoG) and the Hessian detector. Trajectories were formed using the linear assignment problem (LAP) tracker^35^ implemented in TrackMate, and MSD analysis was used for motion parameter estimations on fully detected tracks. The detection accuracy of TrackMate-LoG and SPTnet is similar at low heterogeneous background levels, but the detection accuracy gradually decreases as the heterogeneous background intensity increases. While Hessian detection responds strongly on corners and textured areas, it is more sensitive to heterogeneous backgrounds, leading to the lowest overall detection accuracy (**Fig. 3b, top**). Under the high heterogeneous background conditions (Perlin noise: 40), SPTnet maintained a 0.95 Jaccard index (JI) and outperformed the two TrackMate detectors by 27.4% (Hessian) and 29.6% (LoG) and resulted in at least 32.8% and 17.0% lower errors in estimating the Hurst exponent and generalized diffusion coefficient, respectively (**Fig. 3b, middle and bottom**). Notably, SPTnet maintained track integrity even under highly heterogeneous backgrounds, minimizing missed detections and incorrect track splits (**Fig. 3C and Supplementary Fig. 7**), and the localization accuracy is similar to the two TrackMate detectors with sub-pixel localization enabled (**Supplementary Fig. 6**)

Tracking fast-diffusing molecules with finite camera acquisition time often results in motion blur. To generalize our model for tracking these blurred emission patterns, we augmented the training video by incorporating motion-blurred PSFs (**Fig. 3d**). We then evaluated SPTnet’s performance with increasing generalized diffusion coefficient against two TrackMate options. The detection accuracy declined for all three approaches as the generalized diffusion coefficient increased. SPTnet was more robust against the motion blur, maintaining a JI score of 0.88±0.06, which is consistently higher than TrackMate-LoG (0.70±0.09) and TrackMate-Hessian (0.75±0.07) (**Fig. 3e, top**). Trajectories by two TrackMate detectors yield similar estimation errors on both motion parameters, indicating comparable detection quality on the same subset of motion-blurred tracks (low persistence, less elongated PSFs). SPTnet achieves 0.081±0.016 RMSE on Hurst exponent with consistently lower error compared to TrackMate-LoG (0.126±0.003) and TrackMate-Hessian (0.125±0.003) (**Figure 3e, middle**). The deviation from the theoretical limit at smaller generalized diffusion coefficients can be explained by the localization error, which introduces larger uncertainty on Hurst exponent estimation for trajectories with shorter displacements per frame. For the generalized diffusion coefficient, SPTnet achieves 47.1%±15.6% and 47.4%±15.8% lower RMSE compared to the tested TrackMate methods. (**Figure 3e, bottom**).

### Tracking fluorescent beads in glycerol-water mixtures

We used SPTnet to track freely diffusing fluorescent beads in four different fractions of glycerol-water mixtures within a quasi-2D chamber (**Methods**). The Hurst exponents estimated by SPTnet suggest both 100 nm and 200 nm beads exhibit the Brownian motion under different glycerol concentrations (0.4<H<0.6) (**Fig. 4a, bottom**), as expected for beads in purely viscous fluids^36^. The generalized diffusion coefficient estimated by SPTnet decreases with increasing viscosity, and beads of 100 nm diffuse approximately 2-fold faster than 200 nm beads (1.9±0.3 fold, mean±s.d., average over four different glycerol concentrations, each containing 25 tracks). These estimations are consistent with theoretical predictions based on the Stokes-Einstein equation for Brownian motion particles (**Fig. 4a, top**). For this control experimental condition, SPTnet recovered and accurately distinguished motion dynamics for beads of varied sizes in various glycerol-water mixtures, with a low measurement variance through short 0.6-second videos.

**Figure 4.**
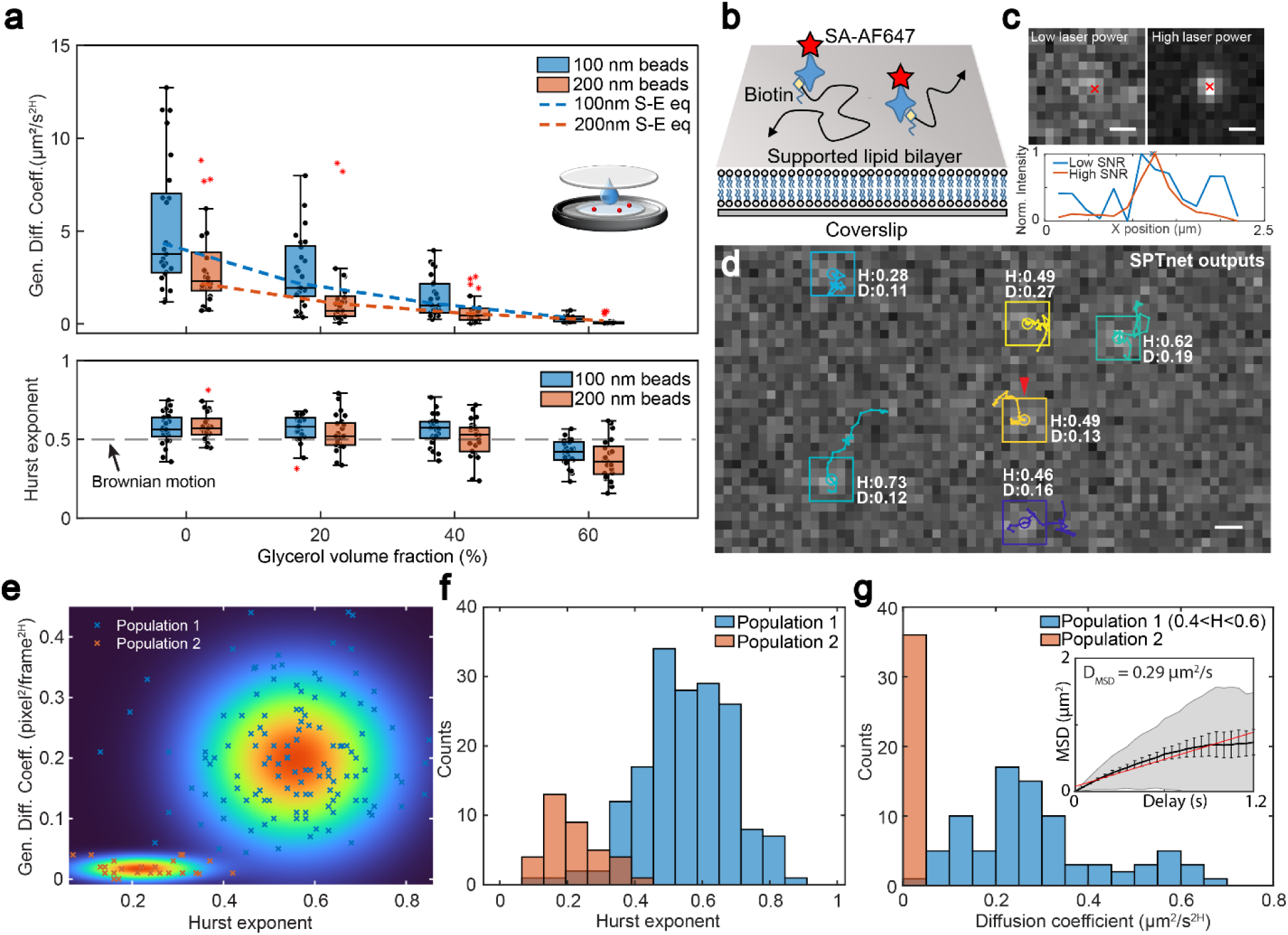
SPTnet tracking of fluorescent beads and fluorescently labeled lipids on SLB. **a,** SPTnet estimated motion parameters for 100 nm and 200 nm fluorescent beads diffusing in water and at different concentrations of glycerol. In the box plot, the center line represents the median, the box limits indicate the 75th and 25th percentiles and the whiskers extend to the most extreme data points that are not considered outliers (red symbols). The dotted lines indicate the theoretical diffusion coefficients predicted from the Stokes-Einstein equation. **b,** Diagram illustrating the tracking of streptavidin-Alexa Fluor 647 (SA-AF647) labeled DOPE-biotin diffusing on supported lipid bilayers (SLBs). **c,** Intensity profiles of PSFs under different excitation laser powers, with SPTnet estimates indicated by red crosses. The low SNR subregion is taken from panel d, marked by the red arrow. **d,** An example of SPTnet outputs. **e,** SPTnet estimated Hurst exponent and generalized diffusion coefficient for all analyzed tracks of AF647-DOPE diffusing on SLBs (n=204), with a 2D Gaussian mixture model overlay distinguishing two populations. **f,** SPTnet estimated Hurst exponent for each identified population. **g,** SPTnet estimated generalized diffusion coefficients, where population 1 includes only motion types close to Brownian motion (0.4<H<0.6). The inset shows diffusion coefficients obtained from the MSD fitting of tracks generated by TrackMate. Error bar: standard error of the mean. Scale bar, 500 nm.

### Tracking diffusion of individual lipids on supported lipid bilayers (SLBs)

SPTnet is capable of analyzing short trajectories under low SNR conditions. We used total internal reflection fluorescence microscopy (TIRF) microscopy and SPTnet to track single lipids, dioleoylphosphatidylethanolamine (DOPE) tagged with Alexa Fluor 647 (AF647) diffusing on SLBs (**Fig. 4b**) – a widely used approach to understand cellular and molecular interactions. AF647 bleached rapidly even under low excitation power, creating a trade-off between SNR and available track length that often challenges conventional algorithms. SPTnet successfully detected PSFs under low SNR conditions without creating spurious detections (**Fig 4c and Fig.4d**). We analyzed 40 subregions, each area measuring 10.2 µm^2^, using SPTnet and generated 204 trajectories (>10 frames) of AF647-DOPE with a mean duration of 0.5±0.1 s (**Fig. 4d, bottom right, and Supplementary Video 3 show example SPTnet outputs**). Among these trajectories, 17.6% were identified as immobile, with a generalized diffusion coefficient of 0.003±0.003 µm^2^/s^2H^ (mean±s.d., N=36) (**Fig. 4e and Fig. 4g**) and a mean Hurst exponent of 0.21±0.08 (**Fig. 4f**). These molecules were trapped in defects in the SLBs, restricting their movement. The other population consists of freely moving molecules exhibiting Brownian motion (H=0.55±0.13, N=168) (**Fig. 4e and Fig. 4f**), with a significantly higher diffusion coefficient of 0.29±0.16 µm^2^/s^2H^ (N=68 for 0.4<H<0.6), consistent with MSD analysis (0.29±0.25 µm^2^/s, N=61) using TrackMate generated tracks (**Fig. 4g**), which require careful tuning of the threshold parameters and exclusion of tracks with low coefficients of determination (R^2^<0.8).

### Tracking Reticulon 4 (Rtn4) and Sec61ꞵ diffusion in live COS-7 cells

Single-particle tracking photoactivated localization microscopy (sptPALM^37^) enables studies of high spatiotemporal motion dynamics in live cells. However, a typical sptPALM dataset contains thousands of single-molecule traces which are often quite short, making data processing time-consuming and error-prone with conventional approaches. Here, we demonstrate SPTnet by tracking two different types of ER proteins and resolving their distinct behaviors. First, we tracked the diffusion of Rtn4, an endoplasmic reticulum (ER) membrane protein that promotes ER membrane curvature^38, 39^, in live COS-7 cells. Individual Rtn4 molecules expressing HaloTag^40^ were labeled with PA-JF_549_-HaloTag ligand^41^, which allows sparse activation and tracking. Using SPTnet, we identified over 4,000 trajectories lasting more than 10 frames, along with their corresponding motion parameters, from 1,000 camera frames over 20 seconds. (**Fig. 5b**).

**Figure 5.**
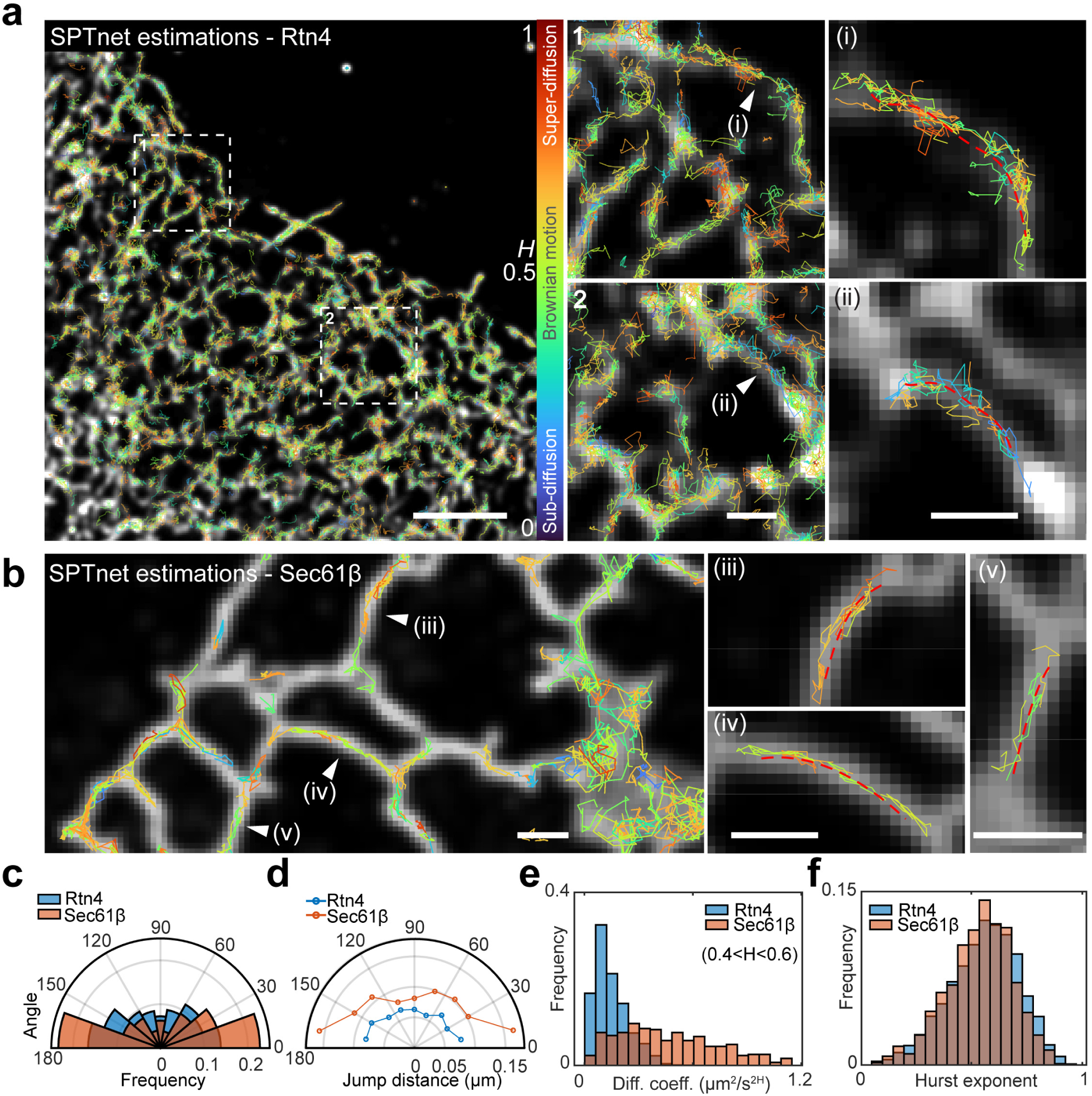
Single particle tracking photoactivated localization microscopy (sptPALM) of Rtn4 and Sec61ꞵ analyzed by SPTnet. **a,** Image of cumulative single-particle tracks (>10 steps) of Rtn4b-HaloTag labeled with PA-JF_549_-HaloTag ligand in live COS-7 cells. Tracks are generated by SPTnet and overlaid with averaged widefield signal. Color coding represents the estimated Hurst exponent. Zoomed-in views of the dotted white box are shown on the right. Panels (i) and (ii) display tracks close to the red dotted line marking the ER tubule centerlines. **b,** SPTnet analyzed sptPALM data of Sec61ꞵ-HaloTag labeled PA-JF_549_-HaloTag ligand, with (iii), (iv), (v) showing tracks using the same methods as Rtn4. **c,** The jump angle distribution for Rtn4 and Sec61ꞵ. For each step along the track, the angle is calculated relative to the nearest position on the ER tubule centerline. **d,** Mean jump distance of Rtn4 and Sec61ꞵ, binned by jump angle of 20°. The jump distance represents the Euclidean distance between two consecutive steps in a track**. e,** Distribution of generalized diffusion coefficients for Brownian-like (0.4<H<0.6) tracks of Rtn4 (n=1829) and Sec61ꞵ (n=434). **f,** Distribution of Hurst exponents for analyzed Rtn4 (n=4463) and Sec61ꞵ trajectories (n=926). Scale bar, 5 µm (**a**) left panel, 1 µm (**a**(2), **a**(ii),(**b**)).

Next, we tracked the ER translocon channel protein Sec61β and compared its motion behavior to Rtn4 (**Fig. 5**). By measuring jump angles relative to the ER tubule centerline (**Supplementary Note 9**), we observed notable differences in jump angle distributions between Rtn4 and Sec61β (**Fig. 5c**). We found 40.7% of Rtn4 jumps (of 760 analyzed jumps) are in oblique or perpendicular directions (45° < angle < 135°) relative to the ER elongation axis. In contrast, Sec61β exhibited a much stronger preference for movement along the axial direction of the ER tubules, with only 26.6% (of 496 analyzed jumps) occurring in oblique or perpendicular directions, consistent with our observed back-and-forth motion of Sec61β along the ER tubules (**Supplementary Video 6 and 8**). Additionally, jump distance analysis revealed that Sec61β exhibits significantly larger step sizes compared to that of Rtn4 (P<0.0001, two-tailed t-test). On average, Sec61β moves 0.14±0.10 µm per step (n=364) along the tubules (< 45° or > 135°) compared to 0.10±0.07 µm (n=132) in other directions (**Fig. 5d**). Rtn4 shows a relatively more uniform distribution of jump distances across angles, with 0.08±0.05 µm (n=451) along the tubules and 0.06±0.05 µm (n=309) in other directions (**Fig. 5d and Supplementary Video 5 and 7**).

Motion behavior analysis of Rtn4 showed a Hurst exponent distribution for overall tracks similar to that of Sec61ꞵ, both being close to Brownian motion with weak persistence (**Fig. 5f**). Among Brownian-like tracks (0.4<H<0.6), Rtn4 diffuses more slowly (P<0.0001, two-tailed t-test), with a generalized diffusion coefficient of 0.15±0.09 µm^2^/s^2H^ (n=1829) compared to Sec61ꞵ of 0.46±0.27 µm^2^/s^2H^ (n=434) (**Fig. 5e**). It is important to note that these measurements are based on the 2D projection of diffusion on a 3D cylindrical structure, which effectively wraps the trajectories and may lead to underestimation of the diffusion rates.

## DISCUSSION

SPTnet is a novel deep learning framework capable of processing videos of single molecules and simultaneously predicting their trajectory and motion parameters with high accuracy. This end-to-end approach simplifies the conventional SPTnet pipeline and enhances accuracy by efficiently incorporating global spatial-temporal information to optimize the trajectory prediction and motion analysis simultaneously. We calculated theoretical precision limits for estimating Hurst exponent and generalized diffusion coefficient across varying lengths of trajectory, motion types, and SNRs. Our results demonstrated that the precision achieved by SPTnet approaches these theoretical information limits. We showed that SPTnet is versatile and can be trained to adapt to different imaging modalities, including TIRF, epi-illumination, and low NA objective (**Supplementary Fig. 5**). It performs robustly even in challenging imaging conditions, such as heterogeneous backgrounds, motion blur, and high noise. We also evaluated the performance of SPTnet through the 2nd AnDi challenge datasets^42^. Under scenarios suitable for SPTnet in this competition, it achieved the lowest errors in estimating the anomalous exponent and generalized diffusion coefficient compared to other methods (**Supplementary Note 10**).

SPTnet is also easy to use by directly taking raw camera frames as input, bypassing the empirical and error-prone process of parameter tuning processes involved in each step of the conventional particle tracking pipeline. Once trained, SPTnet provides instantaneous (<60 ms, for a 30-frame video) trajectory and motion parameter prediction, with its computation time independent of molecule numbers and the duration of fluorescence emission. Training, however, takes approximately 100 hours on a high-end GPU (Nvidia RTX 3080) due to the model’s large parameter set in 3D convolutional layers and transformers. To this end, we implemented an information-weighted loss function to accelerate convergence^27^. Additionally, users can apply transfer learning^43^ to a pre-trained model significantly reducing the initial training time required for the neural network.

In this work, we trained SPTnet primarily on a single anomalous motion model – fractional Brownian motion. This choice is well-suited for live cell imaging on a microscopic level with consecutive frames and short frame rates, whereas the CTRW and LW models often exhibit long stationary waiting periods and ballistic jumps, characteristic of the molecular dynamics of mesoscopic scale^17^. In addition, fBm generalizes various types of motion suitable to distinguish diverse biomolecule motion behaviors with a unified parameter, including superdiffusion (direct motion), Brownian motion, and subdiffusion (confined motion). However, the framework of SPTnet can be adapted to accommodate additional motion models by shifting from binary to multi-class prediction with minor architectural modifications. Although current SPTnet analyzes 30 video frames considering the sufficient estimation precision, it can handle long trajectories with post-processing, for instance, SPTnet can be used in a running window analysis with track linking between different video segments to study motion behavior switching in long trajectories (**Supplementary Video 4**). Given the previous successes in 3D single-molecule localization achieved using deep neural networks^27, 44–46^, we expect SPTnet can be extended to 3D particle tracking. SPTnet framework allows the incorporation of aberrated single-molecule PSFs into training videos, and such design can also facilitate 3D tracking through aberrated specimens, such as whole cells and tissues, with high accuracy.

SPTnet allows direct SPT analysis on videos, reducing the complexity of the current SPT pipeline, and the holistic view efficiently extracts global spatial-temporal information allowing high precision analysis of motion dynamics. The SPTnet framework is flexible with easily customizable code to generate training videos to meet the diverse needs of different users. We believe SPTnet provides an easy-to-use, fast, and accurate deep learning-based SPT analysis framework, especially for conditions that are previously considered challenging.

### Data availability

Example training and testing data for SPTnet are available in supplementary software packages. Complete training and testing datasets can be generated through the shared codes. Other data that support the findings of this study are available from the corresponding authors upon request.

### Code availability

Python scripts for training, validating, and testing SPTnet, as well as MATLAB codes for generating single-molecule training videos are available as Supplementary Software. Further updates will be made available at https://github.com/HuanglabPurdue/SPTnet.

## Author contributions

C.B. and F.H. conceived the project and developed SPTnet. C.B. wrote the software, performed experiments, and analyzed the data. C.B., K.L.S., Y.Z., S.T.L.-N, and F.H. designed the experiments and prepared supported lipid bilayers and biological samples. M.M and Y.Z. aligned and maintained imaging systems. S.T.L.-N and F.H. supervised the study. All authors wrote the manuscript.

## Acknowledgments

We would like to thank Peiyi Zhang and Yilun Li for their helpful discussions on the Fisher information in fractional Brownian motion, and Yuxuan Feng for her suggestions on the notation used in formulas. We thank Hao-Cheng Gao and Li Fang for their contribution to the live-cell imaging system. We thank Thomas D. Pollard for originating the idea of SPTnet and for providing suggestions on the manuscript. We thank Christopher J. Obara and Jennifer Lippincott-Schwartz for suggestions on the ER membrane proteins. We also thank Gorka Muñoz-Gil for his valuable support with the AnDi challenge dataset. This work was supported by the US National Institutes of Health (grants GM119785 to F.H., and GM132024 to K.L.S).

## Methods

### SPTnet architecture

Following the concept of applying transformer for object detection in images^20^, SPTnet consists of sequentially connected components: a 3D-ResNet backbone, the two-stream decoder-encoder transformers with a feature fusion module, and fully connected layers for different output tasks. In the backbone, after an initial 3D convolutional layer, four stages of 3D residual blocks are used, and between each stage we downsample spatial dimension to half and maintain the temporal dimension with maxpooling layers. The number of filters is doubled along with each spatial dimension reduction. Within each residual block, 3×3×3 filters are used for convolution, and ReLU was used as an activation function between convolutional layers. The extracted features are then processed by either incorporating a 2D positional embedding and flattening frame-wise into a 1D vector for input to the Spatial Transformer (Spatial-T) or using a 3D positional embedding and flattening for input to the Temporal Transformer (Temporal-T). We followed the original encoder-decoder transformer design with minor adjustments, eliminating dropout layers and adjusting input dimensions to accommodate sequences of microscopy frames. Both Spatial-T and Temporal-T consist of 6 stacked transformers with 8 multi-attention heads. Detailed architecture and parameters used in SPTnet can be found in Supplementary Figure 3 and Supplementary Table 1.

### Training video simulation

To generate realistic training videos, we simulated a random number of particle motion trajectories using 2D fractional Brownian motion (fBm), with Hurst exponent and generalized diffusion coefficient sampled from uniform distributions. Each particle’s trajectory was assigned a random starting frame and duration within the field of view. Point spread functions (PSFs) were modeled based on scalar diffraction theory, with phase-retrieved Zernike coefficients for simulating aberrated or in-focus PSFs. Heterogeneous backgrounds were introduced using Perlin noise to mimic structured noise from biological structures or out-of-focus emitters. For fast-moving particles, motion blur was incorporated by subdividing exposure times into finer intervals, generating a blurred PSF by summing PSFs across these intervals. (see Supplementary Note 4 for detailed descriptions)

### Training details

We split 200,000 simulated videos randomly into training (80%) and validation (20%) sets. Each video contained 30 frames with width and height of 64×64 pixels. The Input data were normalized based on the maximum and minimal intensity of the entire video before being processed by the neural network. Our model was trained on a single machine equipped with an NVIDIA GeForce GTX3080 GPU. We utilized AdamW^47^ as the optimizer with learning rate set to 0.0001. Additionally, the early stopping mechanism was applied to avoid overfitting, whereby training would stop if the validation loss failed to decrease over 6 consecutive epochs.

### Coverslip cleaning

25-mm-diameter #1.5 coverslips (Thorlabs CG15XH) were soaked in 1 M KOH aq (Fisher P250-1) and ultrasonicated for 15 min using an ultrasonic cleaner (Branson M2800H). Subsequently, the coverslips were washed thoroughly sonicating for 15 min in HPLC water (Fisher W5-4), 200-proof ethanol (Fisher BP28184), and HPLC water one after another. After drying in an oven (Avantor 89508-424) at 110 °C for 1 hr, the coverslips were either sterile under UV for 2 hr for cell samples or directly used for beads samples.

### Beads diffusing in glycerol-water mixtures

In this experiment, we prepared 1:10^7^ dilution of fluorescent beads (FluoSpheres™ Carboxylate-Modified Microspheres, Invitrogen™) on an octadecyltrichlorosilane (Sigma-Aldrich 104817) treated coverslip, following previously described methods^48^ to coat the glass surface with hydrophobic groups and prevent the adsorption of beads on the surface. The sample was illuminated with a 642 nm laser, and beads movement was recorded using an sCMOS camera on a custom-built microscope^24^ with a 0.4 NA objective and 600 nm pixel size. The use of a low NA objective renders the PSF shape less sensitive to aberrations and axial shifts, allowing us to acquire quasi-2D diffusion data in a thin chamber.

### Supported lipid bilayers (SLBs) preparation

SLBs were prepared as previously described^49^. Briefly, coverslips were treated with Piranha solution and then sealed in a metal imaging chamber (ThermoFisher A7816). A lipid vesicle solution (0.5 mg/mL) containing 95 mol% 1,2-Dioleoyl-sn-glycero-3-phosphocholine (DOPC, Avanti 850375), 4 mol% 1,2-dioleoyl-sn-glycero-3-[(N-(5-amino-1-carboxypentyl) iminodiacetic acid) succinyl] (nickel salt) (DGS-NTA(Ni), Avanti 790404), 1 mol% 1,2-dioleoylsn-glycero-3-phosphoethanolamine-N-(cap-biotinyl) (DOPE-biotin, Avanti 870273P) were added to the holder in a 2x PBS solution. After mixing the solution in the chamber and a 15-minute incubation at room temperature, the SLBs were washed three times with 1x PBS. Finally, 200 pM Streptavidin Alexa Fluor 647 conjugate was added to the holder, followed by the same incubation and washing steps with PBS before imaging on a Nikon Ti2 TIRF system.

### Cell culture

COS-7 (ATCC) cells were cultured in Dulbecco’s modified Eagle’s media (DMEM, high glucose, GlutaMAX™ Supplement, Gibco 10566) with 10% v/v fetal bovine serum (FBS, Gibco A5670701) and 100 U/mL penicillin-streptomycin (P-S, Gibco 15070063) in a 5% CO2 humidity incubator at 37 °C.

### Stable cell line generation using Lentivirus

The lentiviral particles were designed and constructed by VectorBuilder Inc. and contain the following vectors: a HaloTag expression vector targeting human Rtn4b (VB230512-1421eyq). COS-7 cells were seeded in 24-well plates at a density of 1.2 × 105 cells per well. After overnight incubation, cells were infected with lentiviral particles expressing an Rtn4-HaloTag according to the manufacturer’s protocol using polybrene transfection reagent and subsequently kept in a 5% CO2 incubator at 37 °C for 48 h. The medium was replaced with fresh complete culture medium, and the cells were cultured for 2 days after transduction. Cells were subjected to blasticidin (Gibco A1113903) selection for an additional 72 h. The single colonies of the transfected cells were further sub-cultured in a 96-well plate and selected by fluorescence microscopy for the homogeneity of expression level.

### Transient mammalian cell transfection

COS-7 cells were overnight cultured on precleaned 25-mm-diameter #1.5 coverslips in phenol red-free DMEM (FluoroBrite™ DMEM, Gibco A1896701) containing 10% FBS (imaging media) at 40% confluency. Plasmid (Halo-Sec61-C-18, Addgene plasmid 123285) transfection was performed using Lipofectamine 3000 (Invitrogen L3000-008) according to the manufacturer’s specifications, using ∼2,600 ng per well in 6-well plates (Corning, CLS3516) and incubating for 36 hrs in the incubator at 37 °C before sptPALM labeling.

### Live cell sptPALM labeling

Cells were overnight cultured on precleaned 25-mm-diameter #1.5 coverslips in the imaging media at 60% confluency and then labeled by incubating with 300 nM PA-JF_549_-HaloTag ligand (generous gift from Dr. Luke D. Lavis, Janelia Research Campus, HHMI) in imaging media for 1 hr at 37°C preventing from light. After washing twice with imaging media, the coverslips were mounted into the Attofluor™ Cell Chamber (Invitrogen A7816), and 2 mL imaging media (prewarmed at 37°C) was added for PALM imaging.

### sptPALM data acquisition

Live cell sptPALM data were taken by a custom-built total internal reflection fluorescence (TIRF) setup, equipped with a 523 nm (CNI MLL-FN-523.5/300 mW), a 560 nm (MPB Communications F-04306-102/1 W) and a 647 nm (MPB Communications F-04306-113/1 W) laser for excitation, a 405 nm (CrystaLaser DL-405-100/100 mW) laser for activation, a sCMOS camera (ORCA-Fusion Digital CMOS, Hamamatsu #C14440-20UP), and an Olympus UPlanApo 60x/1.50 Oil HR objective. Laser power was controlled by an acousto-optic tunable module (AA Optoelectronic AOTFnC-400.650-TN), and the laser light was guided through a polarization-maintaining single-mode fiber (Thorlabs PM-S405-XP). An adjustable mirror was used to switch between wide field (epi), highly inclined and laminated optical sheet (HILO), and total internal reflection fluorescence (TIRF) illumination modes. Emission light was separated from the excitation and activation light with a proper combination of a dichroic mirror and a suppression filter installed in an Olympus U-MF2 filter cube for each excitation laser. For all the sptPALM data, we used a combination of a Chroma T588lpxr dichroic and a Semrock FF01-600/52-25 bandpass filter. Objective and filter cubes were installed into an inverted microscope stand (Olympus IX73). Images were acquired with a 20 ms exposure time, and actual excitation laser powers of 560 nm in the back focal plane using peak power densities of ∼0.5 kW/cm^2^. The PA-JF_549_ labels were photoconverted by a 405 nm laser activation at 0.4 W/cm^2^. The setup components were controlled with custom LabView (2015 32bit) software. During imaging data acquisition, cells were maintained alive at 37 °C and 5% CO2 using a stage-top incubator (Tokai-hit STXG-WSKMX-SET).

