## Supplementary Materials for "SPTnet: a deep learning framework for end-to-end single-particle tracking and motion dynamics analysis"

#### List of Additional Supplementary Materials

#### Supplementary Figures

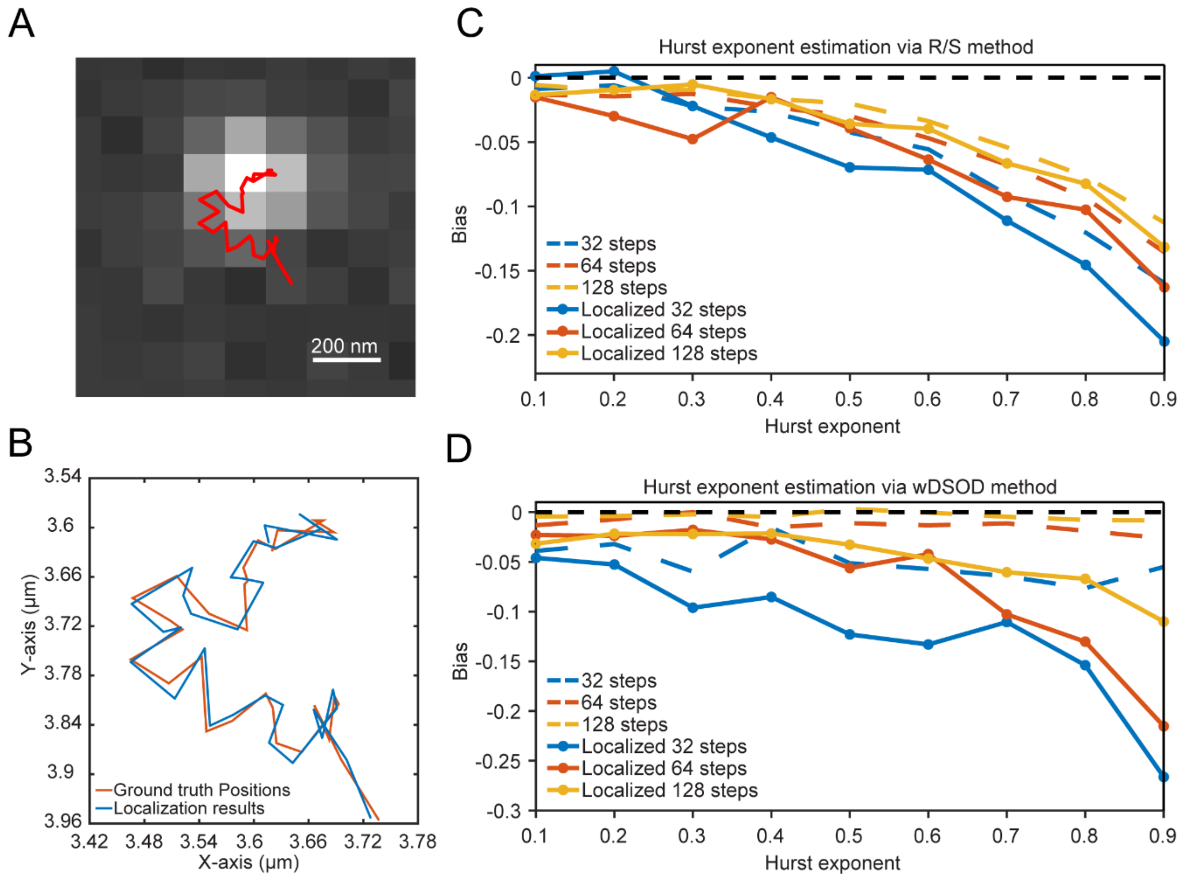

**Supplementary Fig. 1: Localization imprecision and short trajectory lengths introduce bias in Hurst exponent estimation using the R/S and wDSOD methods.** (A) An example subregion of the simulated PSF overlaid with its ground truth trajectory. (B) Localizations of 32-frame image stacks using maximum likelihood estimation (MLE)-based 2D Gaussian fitting (INSPR toolbox1), with results (blue) compared to the ground truth positions (red). (C) Bias in Hurst exponent estimation using 32, 64, and 128 steps of ground truth and localized coordinates. For each Hurst exponent condition, the bias is calculated from 1000 simulated data via the R/S method. (D) Same as (C) but using the wDSOD method for estimation.

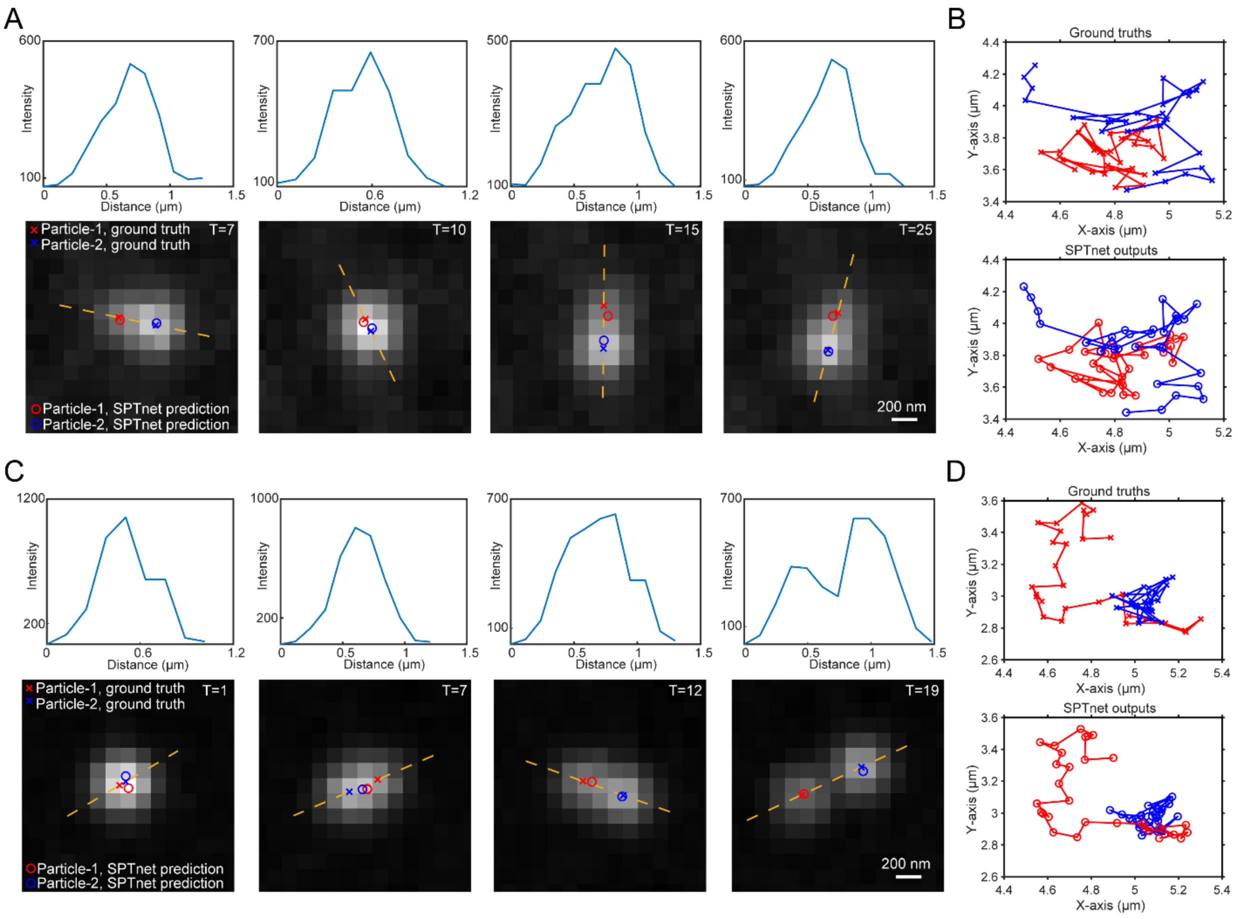

**Supplementary Fig. 2: SPTnet estimations on spatially overlapping emitters.** (**A, C**) Examples of SPTnet estimation results from a simulated video across four selected frames. Each subregion contains two PSFs with ground truth positions marked by red and blue crosses. SPTnet estimations are represented as circles with the same color coding. The intensity profile along the yellow dotted line is shown above each subregion. (**B, D**) The entire 30-step trajectories of two particles from (A) and (C). SPTnet consistently tracked both particles throughout the video without missing detections or incorrect trajectory splitting

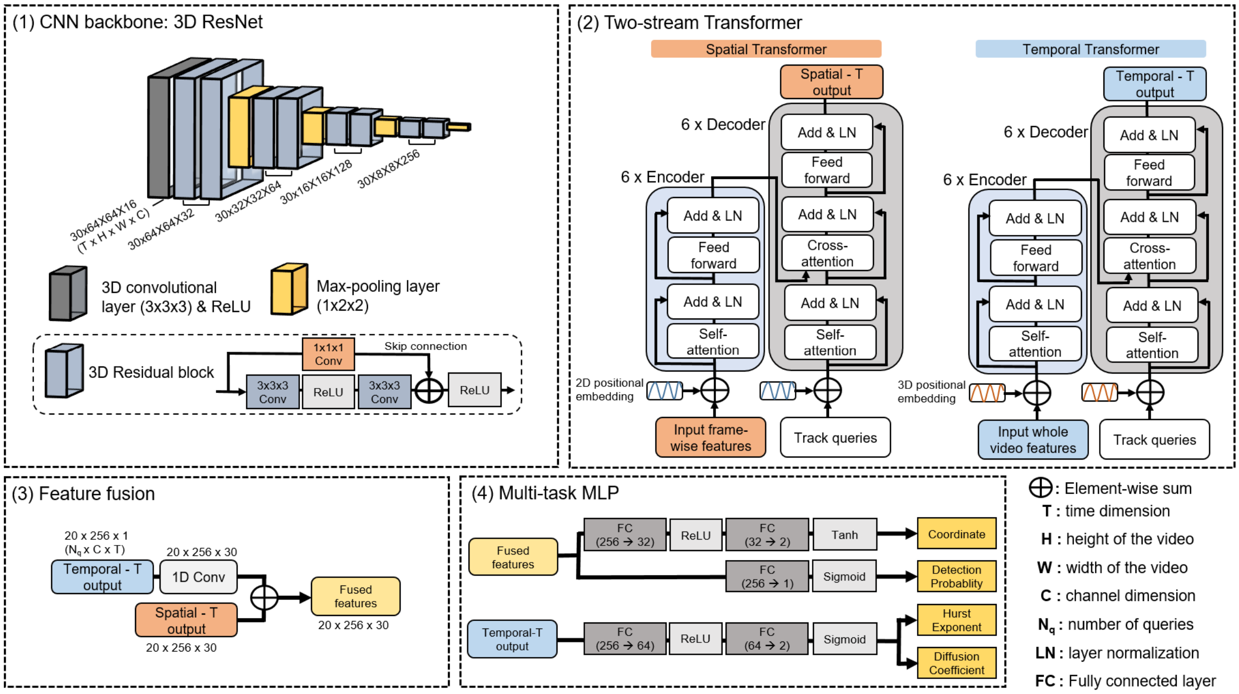

**Supplementary Fig. 3: Detailed architecture of SPTnet.** SPTnet consists of four sequentially linked modules. Input videos are first processed by a modified 3D ResNet, which uses convolutional layers for initial feature extraction. The resulting feature maps are then analyzed by the two-stream transformers module, where Spatial-T processes frame-wise feature maps, while Temporal-T analyzes all features across the entire video. Next, the outputs from the two transformers are fused together and passed to a multilayer perceptron (MLP) to estimate particle presence probabilities and positions at each frame for all track queries. The output from Temporal-T is processed by another MLP to predict the Hurst exponent and generalized diffusion coefficient.

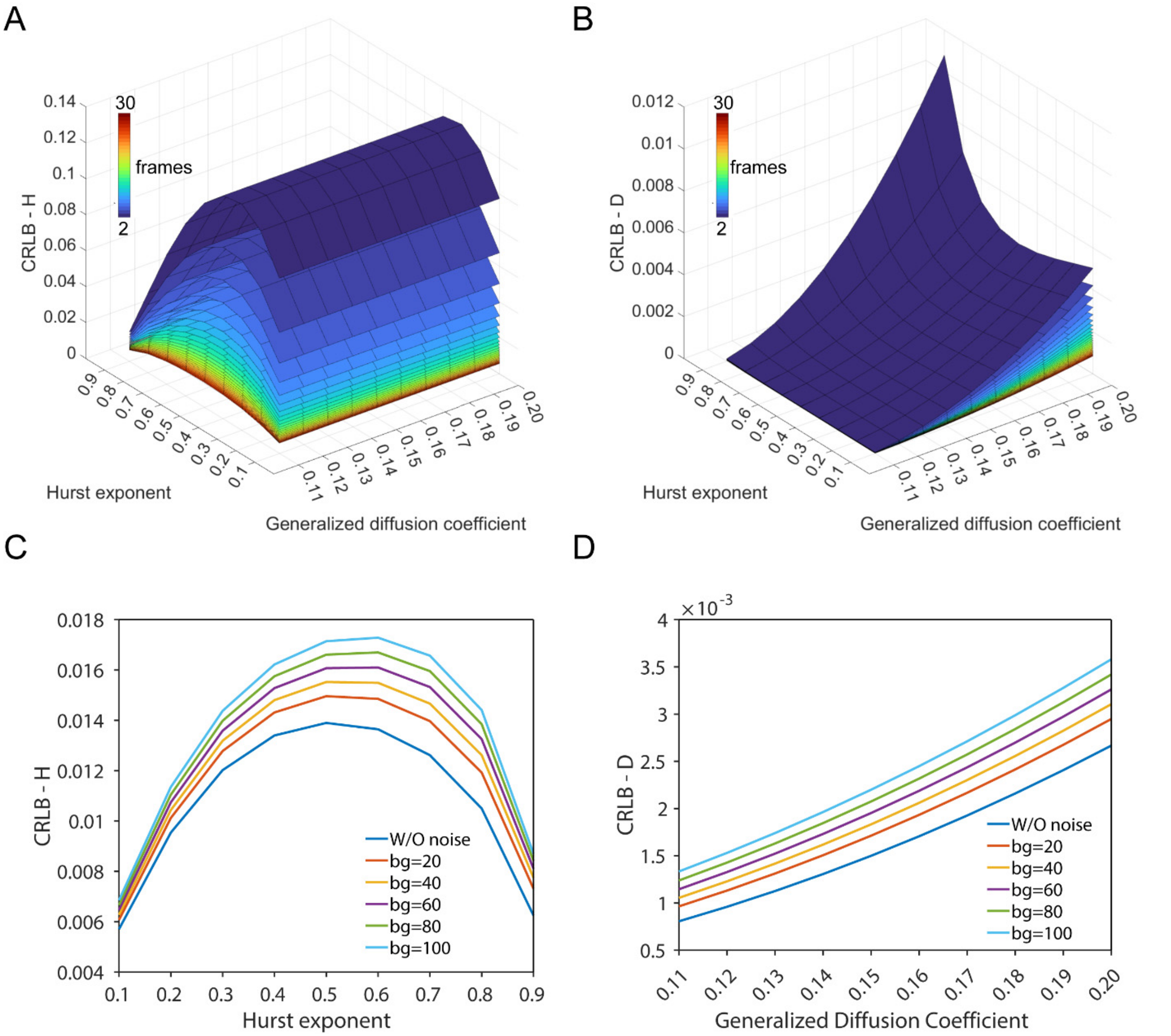

**Supplementary Fig. 4: Cramer-Rao lower bound (CRLB) for estimating different Hurst exponents and generalized diffusion coefficients.** (**A**) Without localization error, the CRLB of Hurst exponent for fBms with different trajectory lengths is represented by different colors. Trajectories with shorter lengths result in higher CRLB values. (**B**) Similar to (A), the CRLB is shown for estimating the generalized diffusion coefficient. (**C**) CRLB for estimating the Hurst exponent from 30 steps fBm trajectories with generalized diffusion coefficient of 0.15 pixel$^2$/frame$^{2H}$ in the presence of different localization errors. The localization errors are based on the CRLB for estimating the positions using a point spread function (PSF) model with 500 photon counts and Poisson noise under varying background levels. (**D**) Similar to (C), but showing the CRLB for estimating the generalized diffusion coefficient with a Hurst exponent of 0.5.

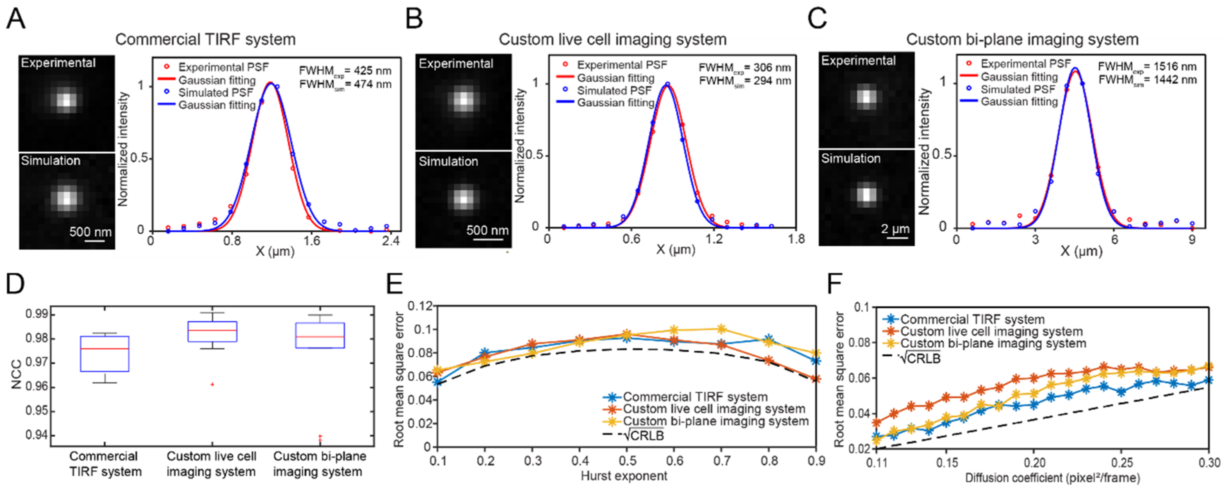

**Supplementary Fig. 5: Evaluation of discrepancies between experimental and simulated PSFs and SPTnet performance across different imaging systems.** (**A**, **B**, **C**) Comparison of experimental PSFs from fluorescent beads with simulated PSFs generated via the pupil function across different imaging systems. The intensity profiles along the x-axis of experimental PSFs are shown in red, while the simulated PSFs are shown in blue. The full width at half maximum (FWHM) of each intensity profile is calculated based on the fitted Gaussian sigma. (**D**) The similarity between measured and simulated PSFs was assessed using normalized cross-correlation (NCC), with n=10. (**E**) SPTnet performance on Hurst exponent estimations, trained for different imaging systems. (**F**) Similar to (E), but demonstrating diffusion coefficient estimations. The square root of the Cramer-Rao lower bound (CRLB) in the plots provides a general reference for the theoretical estimation precision limit for all three imaging systems and does not account for localization errors.

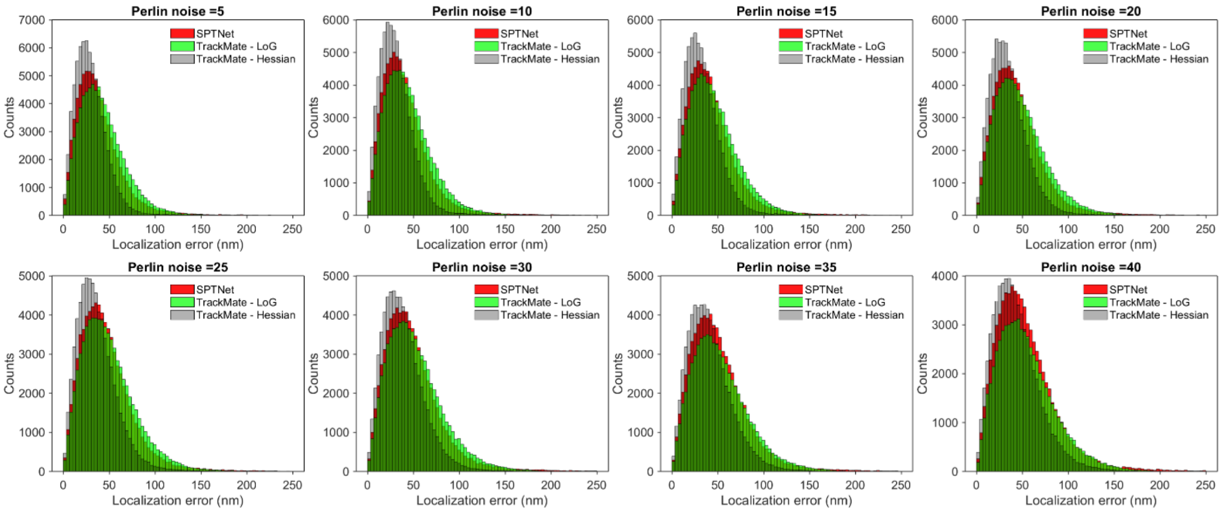

**Supplementary Fig. 6: Localization accuracy of SPTnet compared to TrackMate detectors with subpixel localization under different levels of heterogeneous background.** For each noise condition, we evaluated 1,000 30-frame videos containing a random number of particles. Localization error was calculated as the Euclidean distance between matched particle pairs (see **Supplementary Note 7**), excluding spurious and missed detections. Parameters for each TrackMate detector were manually optimized (see **Supplementary Table 4**).

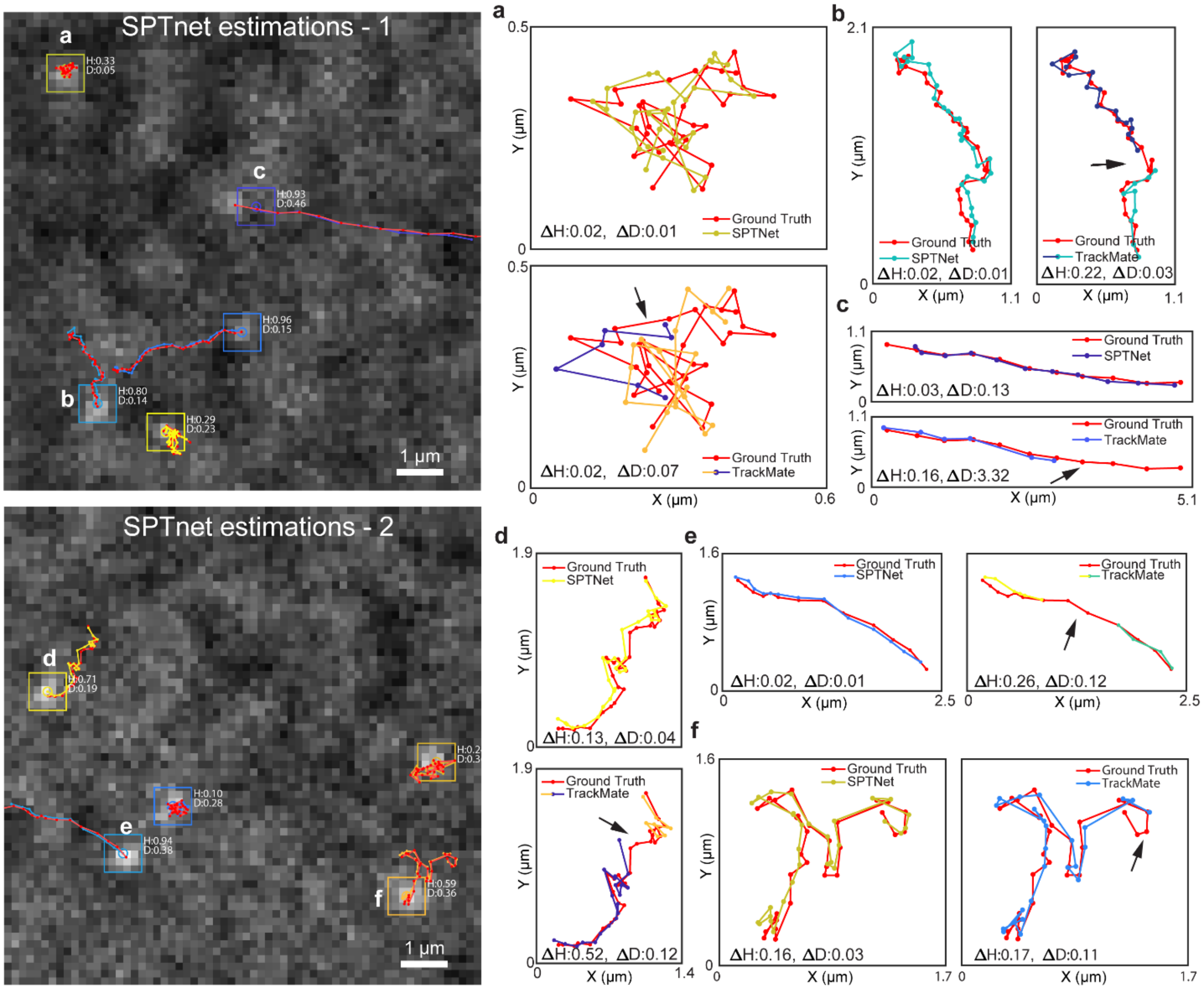

**Supplementary Fig. 7: Comparison of SPTnet and TrackMate estimation results under heterogeneous background conditions.** SPTnet simultaneously estimated the trajectory, Hurst exponent and generalized diffusion coefficient for each moving particle in the input videos. The estimated trajectories from SPTnet are overlaid on the ground truth (shown in red), with boxes indicating the estimated initial positions and "H" and "D" representing the estimated Hurst exponent and generalized diffusion coefficient, respectively. Zoomed-in trajectories **a**, **b,** and **c** from the first video are shown in the corresponding panels on the right, and similarly for **d**, **e,** and **f** from the second video. TrackMate trajectories were generated using a LoG detector and LAP tracker, followed by the parameter estimation for each trajectory using MSD analysis (see **Supplementary Table 4**). Black arrows indicate missing detections and incorrect splitting. For Hurst exponent and generalized diffusion coefficient, the differences between estimations and ground truths are shown as $\Delta H$ and $\Delta D$, respectively.

**Supplementary Table 1. Detailed sizes in each component of the SPTnet architecture**

| Backbone building blocks | Number of building blocks | Components | Kernel/stride/padding | Input dimensions* | Output dimensions* |
|---|---|---|---|---|---|
| Conv - 1 | 1 | -> 3D convolutional layer<br>-> ReLU | [3,3,3],[1,1,1],[1,1,1] | [1,T,H,W] | [16,T,H,W,] |
| Residual Block - 1 | 2 | -> 3D convolutional layer<br>-> ReLU<br>-> 3D convolutional layer<br>-> Short connection<br>-> ReLU | [3,3,3],[1,1,1],[1,1,1]<br>-<br>[3,3,3],[1,1,1],[1,1,1]<br>-<br>- | [16,T,H,W] | [32,T,H,W] |
| Max pooling | 1 | -> 3D Max pooling | [1,2,2],[1,2,2],- | [32,T,H,W] | [32,T,H/2,W/2] |
| Residual Block - 2 | 2 | -> 3D convolutional layer<br>-> ReLU<br>-> 3D convolutional layer<br>-> Short connection<br>-> ReLU | [3,3,3],[1,1,1],[1,1,1]<br>-<br>[3,3,3],[1,1,1],[1,1,1]<br>-<br>- | [32,T,H/2,W/2] | [64,T,H/2,W/2] |
| Max pooling | 1 | -> 3D Max pooling | [1,2,2],[1,2,2],- | [64,T,H/2,W/2] | [64,T,H/4,W/4] |
| Residual Block - 3 | 2 | -> 3D convolutional layer<br>-> ReLU<br>-> 3D convolutional layer<br>-> Short connection<br>-> ReLU | [3,3,3],[1,1,1],[1,1,1]<br>-<br>[3,3,3],[1,1,1],[1,1,1]<br>-<br>- | [64,T,H/4,W/4] | [128,T,H/4,W/4] |
| Max pooling | 1 | -> 3D Max pooling | [1,2,2],[1,2,2],- | [128,T,H/4,W/4] | [128,T,H/8,W/8] |
| Residual Block - 4 | 2 | -> 3D convolutional layer<br>-> ReLU<br>-> 3D convolutional layer<br>-> Short connection<br>-> ReLU | [3,3,3],[1,1,1],[1,1,1]<br>-<br>[3,3,3],[1,1,1],[1,1,1]<br>-<br>- | [128,T,H/8,W/8] | [256,T,H/8,W/8] |
| Max pooling | 1 | -> 3D Max pooling | [1,2,2],[1,2,2],- | [256,T,H/8,W/8] | [256,T,H/16,W/16] |

| Transformers** | Encoder layers | Decoder layers | Number of attention heads | Feedforward dimension | drop out | Normalization before layers | Output dimensions |
|---|---|---|---|---|---|---|---|
| Spatial-T | 6 | 6 | 8 | 1024 | 0 | No | [T,Q,256] |
| Temporal-T | 6 | 6 | 8 | 1024 | 0 | No | [1,Q,256] |

| Multilayer perceptron | Components | Input dimensions | Output dimensions |
|---|---|---|---|
| Coordinate | -> FC 1-1***<br>-> ReLU<br>-> FC 1-2<br>-> Tanh | [T,Q,256]<br>-<br>[T,Q,32]<br>- | [T,Q,32]<br>-<br>[T,Q,2]<br>- |
| Detection probability | -> FC 2<br>-> Sigmoid | [T,Q,256]<br>- | [T,Q,1] |
| Motion parameters (Hurst, Generalized diffusion coefficient | -> FC 3-1<br>-> ReLU<br>-> FC 3-2<br>-> Sigmoid | [Q,256]<br>-<br>[Q,64]<br>- | [Q,64]<br>-<br>[Q,2]<br>- |

* Excluding the batch size dimension. The input and output dimensions in the backbone are channel, frame (T), height (H), and weight (W) of the input videos.

** We use the original transformer modules with minor modifications to the input structure. Q represents the number of track queries, which is set to 20 in most cases.

*** Abbreviation for fully connected layer.

**Supplementary Table 2. Parameters for generating simulation datasets used to train models across different imaging systems**

| Parameters \ Models | | SPTnet models for different imaging systems and application purposes (all models use the same architecture) | | | | |
|---|---|---|---|---|---|---|
| | | Network 1 - Commercial TIRF system | Network 2 – Commercial TIRF system with phase retrieved experimental pupil function | Network 3 – Commercial TIRF system (large diffusion coefficient range for motion blur data) | Network 4 – Custom bi-plane imaging system with Low NA objective | Network 5 – Custom live-cell imaging system with high NA objective |
| Settings of PSF simulation for different Imaging systems | Numerical aperture | 1.49 | 1.49 | 1.49 | 0.4 | 1.50 |
| | Emission wavelength (μm) | 0.69 | 0.69 | 0.69 | 0.69 | 0.58 |
| | Refractive index of the immersion medium | 1.518 | 1.518 | 1.518 | 1 | 1.518 |
| | OTF rescale* | 0.95 | 0.95 | 0.95 | 0.35 | 1.7 |
| | Pixel size (μm) | 0.157 | 0.157 | 0.157 | 0.6 | 0.108 |
| | Aberration (Zernike coefficient)** | - | phase retrieved pupil function | - | - | phase retrieved pupil function |
| Settings for particle dynamics and signal to noise conditions in motion videos | Number of moving particles | $U_i(0, 10)$*** | | | | |
| | Starting time $t_{start}$ (frame) | $U_i(1,30)$ | | | | |
| | Duration (frames) | $U_i(0, 30 - t_{start})$ | | | | |
| | Photon counts per PSF | $U_c(300, 10000)$*** | | | | |
| | Uniform Background | $U_c(1, 50)$ | | | | |
| | Perlin noise background | $U_c(1, 50)$ | | | | |
| | Hurst exponent | $U_c(0.0001, 0.9999)$ | | | | |
| | Generalized Diffusion coefficient (pixel$^2$/frame$^{2H}$) | $U_c(0, 0.5)$ | $U_c(0, 0.5)$ | $U_c(0, 1.5)$ | $U_c(0, 0.5)$ | $U_c(0, 0.5)$ |
| Training and validation data | Number of training data | 160,000 | | | | |
| | Number of validation data | 40,000 | | | | |

* The sigma of the 2D Gaussian filter applied to the optical transfer functions (OTF), which is used to account for the dipole broadening effect.

** Aberrations are only considered in the models used for inferring experimental data. A deformable mirror is used in the custom bi-plane system to compensate for system aberrations.

*** $U_i$ represents a uniformly distributed random integer, and $U_c$ represents a continuous uniformly distributed random number.

**Supplementary Table 3. Simulation parameters used in test datasets**

| Figures | Total number of moving particles | Duration (frames) | Photon counts per PSF | Background intensity | Heterogeneous background intensity | Hurst | Diffusion coefficient (pixel$^2$/frame$^{2H}$) |
|---|---|---|---|---|---|---|---|
| Fig. 2a, left and Supplementary Fig. 5f | 1 | 30 | 1000 | 25 | - | $U_c(0.0001, 0.9999)$ | 0.25 |
| Fig. 2a, right and Supplementary Fig. 5f | 1 | 30 | 1000 | 25 | - | 0.5 | given on the figure |
| Fig. 2b | 1 | 30 | 1000 | 25 | - | given on the figure | given on the figure |
| Fig. 2c | 1 | 30 | given on the figure | given on the figure | - | $U_c(0.0001, 0.9999)$ | $U_c(0, 0.5)$ |
| Fig. 3b and Supplementary Fig. 6 | $U_i(0, 5)$ | 30 | 500 | 10 | given on the figure | $U_c(0.0001, 0.9999)$ | $U_c(0, 0.5)$ |
| Fig. 3c and Supplementary Fig. 7 | $U_i(0, 5)$ | 30 | 500 | 10 | 40 | $U_c(0.0001, 0.9999)$ | $U_c(0, 0.5)$ |
| Fig. 3e | $U_i(0, 5)$ | 30 | $U_i(300, 10000)$ | $U_c(1, 50)$ | $U_c(1, 50)$ | $U_c(0.0001, 0.9999)$ | given on the figure |
| Fig. SS5 | $U_i(0, 10)$ | $U_i(1, 30)$ | $U_i(300, 10000)$ | $U_c(1, 50)$ | $U_c(1, 50)$ | $U_c(0.0001, 0.9999)$ | $U_c(0, 0.5)$ |

**Supplementary Table 4.1: TrackMate settings for heterogeneous background videos**

| Method - 1 | TrackMate Detector: LoG | | | | TrackMate Tracker: LAP tracker | | | | MSD analysis | |
|---|---|---|---|---|---|---|---|---|---|---|
| Testing conditions (heterogeneous backgrounds) | Estimated object diameter (Pixel) | Quality threshold | Pre-process with a median filter | Sub-pixel localization | Max frame to frame linking distance (Pixel) | Gap closing | Gap closing max distance (Pixel) | Max frame gap | Fitting the time-averaged MSD (TAMSD) | Max tau used for fitting (30 frames in total) |
| Perlin bg intensity: 5 | 2.5 | 25 | False | True | 5 | True | 10 | 2 | Least squares | 5 |
| Perlin bg intensity: 10 | 2.5 | 27 | False | True | 5 | True | 10 | 2 | Least squares | 5 |
| Perlin bg intensity: 15 | 2.5 | 29 | False | True | 5 | True | 10 | 2 | Least squares | 5 |
| Perlin bg intensity: 20 | 2.5 | 31 | False | True | 5 | True | 10 | 2 | Least squares | 5 |
| Perlin bg intensity: 25 | 2.5 | 31 | False | True | 5 | True | 10 | 2 | Least squares | 5 |
| Perlin bg intensity: 30 | 2.5 | 33 | False | True | 5 | True | 10 | 2 | Least squares | 5 |
| Perlin bg intensity: 35 | 2.5 | 40 | False | True | 5 | True | 10 | 2 | Least squares | 5 |
| Perlin bg intensity: 40 | 2.5 | 42 | False | True | 5 | True | 10 | 2 | Least squares | 5 |
| Method - 2 | TrackMate Detector: Hessian | | | | TrackMate Tracker: LAP tracker | | | | MSD analysis | |
| Testing conditions (heterogeneous backgrounds) | Estimated object diameter in X & Y (Pixel) | Quality threshold | Normalize quality values | Sub-pixel localization | Max frame to frame linking distance (Pixel) | Gap closing | Gap closing max distance (Pixel) | Max frame gap | Fitting the time-averaged MSD (TAMSD) | Max tau used for fitting (30 frames in total) |
| Perlin bg intensity: 5 - 40 | 2.5 | 0.6 | True | True | 5 | True | 10 | 2 | Least squares | 5 |

**Supplementary Table 4.2: TrackMate settings for motion-blurred videos**

| Method – 1 | TrackMate Detector: LoG | | | | TrackMate Tracker: LAP tracker | | | | MSD analysis | |
|---|---|---|---|---|---|---|---|---|---|---|
| Testing conditions (motion blur) | Estimated object diameter (Pixel) | Quality threshold | Pre-process with a median filter | Sub-pixel localization | Max frame to frame linking distance (Pixel) | Gap closing | Gap closing max distance (Pixel) | Max frame gap | Fitting the time-averaged MSD (TAMSD) | Max tau used for fitting (30 frames in total) |
| Diff. Coeff.*: 0.1 – 1.5 | 4 | 60.26 | False | True | 5 | True | 10 | 3 | Least squares | 5 |
| Method - 2 | TrackMate Detector: Hessian | | | | TrackMate Tracker: LAP tracker | | | | MSD analysis | |
| Testing conditions (motion blur) | Estimated object diameter in X & Y (Pixel) | Quality threshold | Normalize quality values | Sub-pixel localization | Max frame to frame linking distance (Pixel) | Gap closing | Gap closing max distance (Pixel) | Max frame gap | Fitting the time-averaged MSD (TAMSD) | Max tau used for fitting (30 frames in total) |
| Diff. Coeff.*: 0.1 – 1.5 | 3 | 165 | False | True | 5 | True | 10 | 3 | Least squares | 5 |

* Diff. Coeff. : Generalized diffusion coefficient (pixel$^2$/frame$^{2H}$), H: Hurst exponent, and pixel size: 157 nm.

#### Supplementary Note

### 1. Architecture consideration

##### 1.1 Use of 3D-ResNet as backbone

Moving single molecules can overlap with each other, making it challenging to determine the number of emitters and their precise locations from a single frame, especially when the frame is further corrupted by various sources of noise. In such cases, analyzing isolated frames using 2D convolutional layers results in less informative features that rely solely on spatial content, complicating track formation and motion parameter estimation. By extending the standard 2D convolution into 3D, 3D-ResNet[2] incorporates the time dimension, allowing the model to capture temporal relationships across consecutive frames. Additionally, the use of residual connections in 3D-ResNet mitigates information loss and the vanishing gradient problem, enhancing the model's ability to learn complex spatiotemporal features.

The 3D convolutional kernel is set to 3 x 3 x 3 (time, height, width) based on empirical performance from a previous investigation[3]. We eliminated the temporal dimension reduction in the max pooling layers by using a kernel of 1 x 2 x 2 to maintain temporal resolution. The consistent temporal dimension ensures that features align with the per-frame output structure of the final detection and localization. Considering the large memory consumption for 3-dimensional inputs, the batch size used for training on a standard GPU is relatively small (<24), and the variance of the batch statistics can fluctuate significantly between batches. To stabilize the training process, we removed 3D batch normalization between the 3D convolutional layers.

##### 1.2 Use of the two-stream encoder-decoder Transformers

The encoder-decoder architecture originated in the field of machine translation, where it was first introduced to address the challenges of translating long sequences between languages. This architecture functions by encoding input data into a compressed representation and subsequently decoding this representation into the desired output. In SPTnet, the encoder-decoder framework plays a crucial role in achieving precise and robust particle tracking. The encoder extracts spatiotemporal features from the input video, summarizing essential information regarding particle motion and the environmental context. The decoder then utilizes this compressed representation to reconstruct accurate particle trajectories, ensuring consistent tracking across frames.

Transformer, first introduced by Vaswani et al. in 2017[4], is a deep learning architecture designed to address the limitations of recurrent neural networks (RNNs), such as difficulties in modeling long-range dependencies and inefficiencies of sequential data processing. Transformer is able to process all sequence data in parallel through the attention mechanism and compute the weighted sum of the input features based on the $Q$ (query), $K$ (key), and $V$ (value):

$$\mathrm{Attention}(\mathrm{Q},\mathrm{K},\mathrm{V}) = softmax\left(\frac{QK^T}{\sqrt{d_k}}\right)V \tag{1}$$

$\sqrt{d_k}$ is a scaling factor based on the dimension of the key, and the softmax function normalizes the output to produce weighting factors between 0 and 1. In SPTnet, we refer to those queries as track queries. These are embedded vectors representing individual particle trajectories of interest throughout the video. Each track query is designed to estimate the position and motion parameters of a specific particle across multiple frames. These track queries are fed into the transformer decoder, where they interact with encoded spatiotemporal features ($K$ and $V$) through the attention mechanism. This allows each track query to focus on the most relevant features across frames, enabling accurate particle tracking. Initially, we implemented a single transformer to process features from the entire video. While the model accurately estimated motion parameters, each track query could only roughly follow the trajectory and was unable to achieve the sub-pixel-level localization accuracy that conventional detectors can provide. To address this, we refined the model by incorporating another transformer that focuses on frame-wise feature association, significantly improving tracking precision.

The two-stream encoder-decoder transformers with the feature fusion module allow SPTnet to efficiently incorporate molecular motion behavior together with global spatial-temporal information to further refine trajectory reconstruction, thereby overcoming limitations for conventional methods especially when dealing with short trajectories, limited SNR, complex object interactions, and PSF overlapping.

##### 1.3 2D and 3D positional encoding

Transformers do not inherently preserve positional information of inputs (permutation-invariant). Positional encoding, which assigns a unique representation to each position of the input features, is crucial for the model to understand the correlations between pixels and frames. For the inputs to Spatial-T, we encode spatial dimensions of the frame-wise features using 2D sinusoidal positional encoding, following the implementation of the original transformers[4]. On the other hand, the inputs to Temporal-T are encoded across both spatial and temporal dimensions using 3D sinusoidal positional encoding, an extension of the 2D version[5], which can be expressed as,

$$\mathrm{PE(x,y,t,c)} = \begin{cases} \sin\left(\dfrac{x}{10000^{\frac{6i}{D}}}\right), & if\ c = 2i \\ \cos\left(\dfrac{x}{10000^{\frac{6i}{D}}}\right), & if\ c = 2i+1 \\ \sin\left(\dfrac{y}{10000^{\frac{6j}{D}}}\right), & if\ c = 2j \\ \cos\left(\dfrac{y}{10000^{\frac{6j}{D}}}\right), & if\ c = 2j+1+\dfrac{D}{3} \\ \sin\left(\dfrac{t}{10000^{\frac{6k}{D}}}\right), & if\ c = 2k+\dfrac{2D}{3} \\ \cos\left(\dfrac{t}{10000^{\frac{6k}{D}}}\right), & if\ c = 2k+1+\dfrac{2D}{3} \end{cases}, \tag{2}$$

where $x, y, t$ are the width, height, and time dimensions of the inputs, and *i, j,* and $k$ are indices in the range $[0,\ \frac{D}{6})$ for each respective dimension. *D* is the total size of the channel dimension (*c*).

#### 2. Fractional Brownian motion as an anomalous diffusion model

##### 2.1 Fractional Brownian motion

Let $B_H(t)$ denote the fractional Brownian motion (fBm) with a Hurst exponent *H,* defined for time $t \geq 0$. The fBm process $B_H = \{B_H(t)\}$ can be used to model anomalous diffusion, where the relationship between the mean square displacement (MSD) and time is non-linear. In the 1D case, the second moment of fBm is given by,

$$\langle B_H(t)^2 \rangle = 2D_H t^{2H}, \tag{3}$$

where $B_H(t)$ represents the position of a particle undergoing fBm at time *t*, $D_H$ denotes the generalized diffusion coefficient with units of $\mathrm{length}^2 \cdot time^{-2H}$, and *H* is the Hurst exponent. The anomalous exponent α is related to *H* by $\alpha = 2H$.

The Hurst exponent can vary between $0 < H < 1$, regulating the correlation between a moving particle's position at different time points. By adjusting the Hurst exponent, fBm can be linked to the underlying physical mechanisms governing particle diffusion in biological systems[6-8]. For example, when $0 < H < 0.5$, the increments of the motion are negatively correlated, and the particle exhibits sub-diffusion behavior, which can be interpreted as a consequence of particle trapping or transient binding to cellular structures. When $H = 0.5$, the increments are uncorrelated, corresponding to classical Brownian motion. In the case where $0.5 < H < 1$, the increments are positively correlated, resulting in super-diffusion behavior that may represent active transport via motor proteins (**Fig. SS2**).

Modeling anomalous diffusion using the fBm framework offers several advantages. Many biological systems exhibit non-Markovian behavior[9-11], meaning the future dynamics of a biomolecule are influenced by its past, rather than just its present state. The fBm model is well suited to describe this non-Markovian nature due to the memory property of its correlation function. Another advantage of fBm is its self-similarity, which ensures fBm maintains consistent statistical properties across different time scales, making it useful for modeling molecular dynamics that occur over a wide range of scales. Furthermore, fBm provides a succinct and flexible model to describe different motion types in a unified manner through a single parameter, the Hurst exponent.

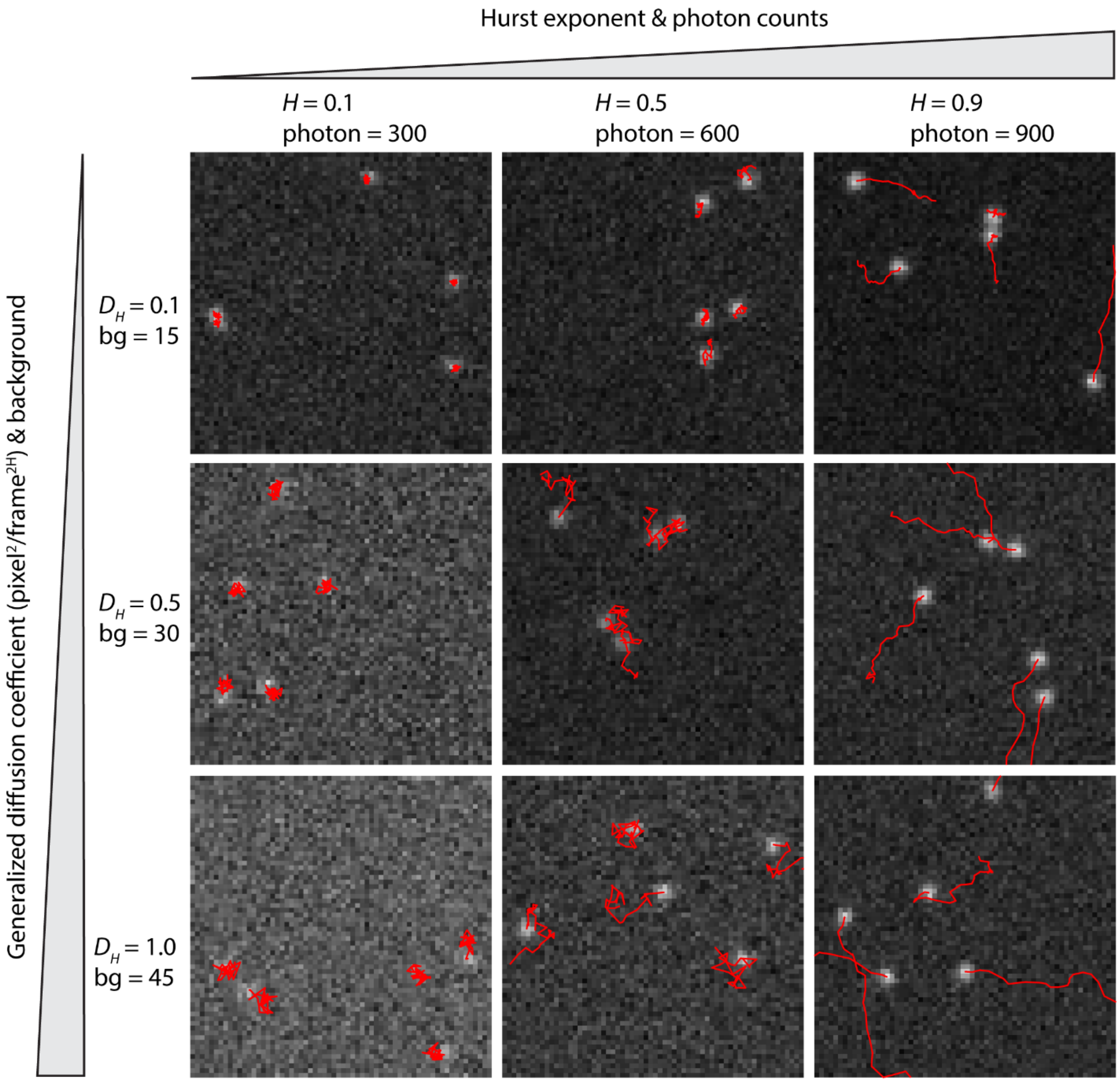

**Fig. SS2: Examples of simulated fractional Brownian motion videos under various Hurst exponents, generalized diffusion coefficients, photon counts, and background conditions**. Each simulated video contains 5 moving emitters and spans for 30 frames, with a uniform background and added Poisson noise. The corresponding trajectories of each emitter are shown in red. The PSFs were simulated based on the parameters of a commercial TIRF microscope (**Supplementary Table 2**)

##### 2.2 Simulation of Fractional Brownian motion sample path

We simulated fractional Brownian motion (fBm) using the Cholesky decomposition method. This approach is straightforward to implement and allows the direct generation of the fBm sample path, without approximating it via cumulative sums of fractional Gaussian noise (fGn), thus providing a more precise representation of the process[12].

To generate a *N* steps sample path of a discrete-time 1D fBm process $B_H = \{B_H(t_i), i = 1, \dots, N\}$

for a given Hurst exponent H and generalized diffusion coefficient $D_H$, we first calculate its covariance matrix $C(t_i, t_j)$,

$$C(t_i, t_j) = E[B_H(t_i)B_H(t_j)] = D_H\left(t_i^{2H} + t_j^{2H} - |t_i - t_j|^{2H}\right), \quad i, j = 1, \dots, N; \tag{4}$$

where $t_i$, $t_j$ are two time points, and $C(t_i, t_j)$ is a symmetric positive definite matrix, allowing it to be decomposed into a unique lower triangular matrix L using Cholesky decomposition,

$$C(t_i, t_j) = \begin{pmatrix} C(t_1, t_1) & \cdots & C(t_1, t_N) \\ \vdots & \ddots & \vdots \\ C(t_N, t_1) & \cdots & C(t_N, t_N) \end{pmatrix} = \mathrm{LL}^T, \tag{5}$$

then the *N* steps fBm sample path can be represented as $X_N = \mathrm{LZ}$, where $\mathrm{Z} = (Z_1, \dots, Z_N)^T$ is a vector of n number drawn independently from a standard Gaussian distribution. The 2D fBm is constructed by combining two independent 1D fBm along the x- and y-axes.

#### 3. Calculation of Cramer-Rao lower bound (CRLB) for estimators in fractional Brownian motion

##### 3.1 CRLB for estimating Hurst exponent and generalized diffusion coefficient

For an unbiased estimator, the best precision it can achieve is limited by the Cramer-Rao lower bound (CRLB). To evaluate the performance of different methods for estimating the Hurst exponent and generalized diffusion coefficient in fractional Brownian motion model, we quantified theoretical precision limits predicted by the CRLB under different motion types and track lengths.

The fBm process of *N* time points (frames), $B_H = \{B_H(t), t = 1, \dots, N\}$ is a centered Gaussian stochastic process characterized by its covariance function $C(\theta)$ (1), where $\theta = (H, D_H)$. The probability density function (PDF) of this multivariate Gaussian distribution is defined as,

$$P(B_H;\ C(\theta)) = \frac{1}{(2\pi)^{\frac{N}{2}} \det[C(\theta)]^{\frac{1}{2}}} exp^{-\frac{1}{2}(B_H{}^T C(\theta)^{-1} B_H)}, \tag{6}$$

we can then base on the likelihood function $L(\theta|B_H)$, which takes the same form as the PDF, to calculate the Fisher information matrix of $\theta$ for the 1D fBm process,

$$\mathrm{I}(\theta)_{ij} = E\left[\frac{\partial \ln L(\theta|B_H)}{\partial \theta_i} \frac{\partial \ln L(\theta|B_H)}{\partial \theta_j}\right],$$
$$\mathrm{I}(\theta)_{ij} = \frac{1}{2} * tr\left(C(\theta)^{-1} \frac{\partial C(\theta)}{\partial \theta_i} C(\theta)^{-1} \frac{\partial C(\theta)}{\partial \theta_j}\right), \tag{7}$$

where $tr$ represents the trace of a matrix. $\mathrm{I}(\theta)_{11}, \mathrm{I}(\theta)_{22}$ are the diagonal terms of the Fisher information matrix of Hurst exponent and generalized diffusion coefficient, respectively. For 2D fBm, since it is constructed from two independent fBm processes $B_H^x$ and $B_H^y$, the fisher information is the sum of the two processes,

$$P_{(X,Y)}\left(B_H^x, B_H^y; C(\theta)\right) = P_X(B_H^x;\ C(\theta)) \cdot P_Y(B_H^y;\ C(\theta)),$$
$$I_{XY}(\theta)_{ij} = E\left[\frac{\partial \ln L(\theta|\mathrm{B}_H^x)}{\partial \theta_i} \frac{\partial \ln L(\theta|\mathrm{B}_H^x)}{\partial \theta_j} + \frac{\partial \ln L\left(\theta|\mathrm{B}_H^y\right)}{\partial \theta_i} \frac{\partial \ln L\left(\theta|\mathrm{B}_H^y\right)}{\partial \theta_j}\right],$$
$$I_{XY}(\theta)_{ij} = tr\left(C(\theta)^{-1} \frac{\partial C(\theta)}{\partial \theta_i} C(\theta)^{-1} \frac{\partial C(\theta)}{\partial \theta_j}\right), \tag{8}$$

then the variance of any unbiased estimator for Hurst exponent $\hat{\theta}_H$ and generalized diffusion coefficient $\hat{\theta}_{D_H}$ is bounded by,

$$\mathrm{Var}\left(\hat{\theta}_H\right) \geq (I_{XY}(\theta))_{11}^{-1},$$
$$\mathrm{Var}\left(\hat{\theta}_{D_H}\right) \geq (I_{XY}(\theta))_{22}^{-1}. \tag{9}$$

##### 3.2 CRLB considering Poisson noise induced localization imprecision

Experimental SPT data is often corrupted by different sources of noise, with the major one being shot noise, which can be modeled through Poisson process. To account for this, we incorporated the influence of Poisson photon statistics into the theoretical precision calculation of motion parameters. Let $X_H(t)$ denote the estimated 1D fBm position at time *t*, including localization imprecision induced by Poisson noise,

$$X_H(t) = B_H(t) + W(t), \tag{10}$$

where $B_H(t)$ represents the ground truth fBm position at time *t*, and $W(t)$ is the corresponding localization error. We assume the localization error follows a Gaussian distribution with a mean equal to the ground truth location and variance given by the CRLB for localization under Poisson noise. The resulting Gaussian random process X = $\{X_H(t_i), i = 1, \dots, N\}$ is characterized by the covariance matrix $C_P(t_i, t_j)$,

$$\begin{aligned} C_P(t_i, t_j) &= E\big[(X_H(t_i) - E[X_H(t_i)])(X_H(t_j) - E[X_H(t_j)])\big], \\ &= \mathrm{E}\big[B_H(t_i)B_H(t_j)\big] + \mathrm{E}\big[\mathrm{W}(t_i)\mathrm{W}(t_j)\big]. \end{aligned} \tag{11}$$

Since the fBm position is independent of the localization error, and localization errors at different time points are also independent, the covariance matrix simplifies to,

$$C_P(t_i, t_j) = \begin{cases} D_H\big(|t_i|^{2H} + |t_j|^{2H} - |t_i - t_j|^{2H}\big) + \sigma_w^2, & i = j \\ D_H\big(|t_i|^{2H} + |t_j|^{2H} - |t_i - t_j|^{2H}\big), & i \neq j \end{cases}, \tag{12}$$

where $\sigma_w^2$ represents the variance of localization error, and it can be calculated by the Fisher information of the likelihood function given the data D follows Poisson process for parameters $\theta$, which include the x, y positions, the total photon counts I, and the background bg. The likelihood function is,

$$L(\theta|D) = \prod_q \frac{{\mu_q}^{D_q} e^{-\mu_q}}{D_q!}, \tag{13}$$

where $\mu_q$ represents the intensity at pixel index *q* given the PSF model. The PSF model used here does not have an analytical expression and was calculated numerically. We assume any two different pixels are independent, and the Fisher information matrix is then given by,

$$I_{loc}(\mathrm{i}, \mathrm{j}) = \sum_q \frac{1}{\mu_q} \frac{\Delta\mu_q}{\Delta\theta_i} \frac{\Delta\mu_q}{\Delta\theta_j}. \tag{14}$$

The variance of the localization error is the CRLB of the *x, y* position estimation as previously defined,

$$\sigma_w^2 = I_{loc}(\mathrm{i}_{xy}, \mathrm{i}_{xy})^{-1}, \tag{15}$$

where $I_{loc}(\mathrm{i}_{xy}, \mathrm{i}_{xy})^{-1}$ denotes the diagonal elements of the inverse Fisher information matrix related to the *x*, *y* positions. The final theoretical estimation precision for *H* and $D_H$ with Poisson noise induced localization imprecision can be calculated through $C_P(t_i, t_j)$ using equation (2). The CRLB of the localization was calculated using the PSF toolbox provided in Supplementary Software.

#### 4. Simulation of the training video

##### 4.1 Particle motion trajectories

We modeled particle trajectories using two-dimensional fBm, characterized by the Hurst exponent $H$ and generalized diffusion coefficient $D_H$. For each particle, $H$ and $D_H$ were randomly sampled from two independent uniform distributions. Each particle was assigned a random starting frame and a trajectory duration of $N$ frames. The particle's trajectory, following a centered 2D-fBm over $N$ frames, was generated using the Cholesky decomposition method (as detailed in the **'Simulation of Fractional Brownian motion sample path'** section). A uniformly distributed random shift ($\Delta x$, $\Delta y$) was then applied to the trajectory to allow for different initial positions of the particles. Class labels were generated for each frame of the trajectories. Trajectories at frames that fell within the particle's duration and inside the field of view (FOV) were labeled as "particle" class, while all other frames were labeled as "background" class. Each track has its own ground truth labels of Hurst exponent $H$, generalized diffusion coefficient $D_H$, coordinate at each frame ($x_t$, $y_t$), and a binary class label per frame.

##### 4.2 PSF generation

Next, we simulated point spread functions (PSFs) at each frame of the previously generated fBm trajectories. PSFs were simulated based on the scalar diffraction theory[13] through the Fourier transform of the pupil function. The pupil function can be generated through phase retrieval as previously described[14]. Briefly, a stack of experimental PSFs at different axial positions ranging from -1 to 1 µm with 100 nm step size was acquired. The noise-reduced experimental data was then processed iteratively using the Gerchberg-Saxton algorithm[15, 16] to generate the pupil function. Finally, the obtained pupil function will be decomposed into Zernike polynomials[17] (Wyant ordering). The PSF position $(x_t, y_t, z_t)$ at frame $t$ was simulated by the Zernike expansion of the pupil function can be expressed as follows,

$$\mu_0(x_t, y_t, z_t) = \left|\mathcal{F}^{-1}\left[h(k_x, k_y)e^{i2\pi k_z z_t}\right]\right|^2, \tag{16}$$

where $h(k_x, k_y)$ is the pupil function, $\mathcal{F}$ is the Fourier transform operator, $k_z = \sqrt{(\frac{n}{\lambda})^2 - k_x^2 - k_y^2}$, $n$ is the refractive index of the objective immersion medium, and $\lambda$ is the emission wavelength. The term $e^{i2\pi k_z z_t}$ represents the defocus and was set to 1, as we assumed the tracked particles undergo 2D diffusion and generate in-focus PSFs ($z_t = 0$). The pupil function can be expressed as,

$$h(k_x, k_y) = \mathrm{A}(k_x, k_y) \cdot \mathrm{e}^{\mathrm{i}\Phi(k_x, k_y)}, \tag{17}$$

where $\mathrm{A}(k_x, k_y)$ and $\Phi(k_x, k_y)$ represent the magnitude and phase of the electric field at the back focal plane of the objective, respectively. The wavefront distortion introduced by the optical system and specimen can be described by a linear combination of a series of Zernike modes ($Z_n$) with amplitude coefficients ($c_n$),

$$\Phi(k_x, k_y) = \sum_{n=1}^{N} c_n Z_n(k_x, k_y) \tag{18}$$

To generalize SPTnet for imaging systems with aberrated PSFs, we used phase-retrieved Zernike coefficients to simulate the training datasets. For models applied to aberration-free systems or simulation data, we set all $c_n$ to zero, assuming a flat wavefront.

The PSFs were normalized and scaled by the total photon count of each corresponding particle. It is important to ensure that SPTnet is capable of dealing with varying signal-to-noise ratio (SNR) conditions, we therefore augmented training datasets by incorporating both uniform ($bg_{unif}$) and heterogeneous ($bg_{structural}$) backgrounds into the simulation. The simulated video at frame *t* can be expressed as,

$$\mu_t(x_{t_k}, y_{t_k}) = \sum_{k=1}^{K} I_k \mu_{0_k}(x_{t_k}, y_{t_k}) + bg_{unif} + bg_{structural}, \tag{19}$$

where $\mu_t(x_{t_k}, y_{t_k})$ is the ideal recorded SPT frame containing a total of *K* PSFs, and $I_k$ is the photon count of the PSF with index *k*. The final step was to corrupt $\mu_t(x_{t_k}, y_{t_k})$ with Poisson noise.

##### 4.3 Heterogeneous background

We use Perlin noise[18] to simulate the potential heterogeneous backgrounds in real microscopy images, which can be introduced by out-of-focus emitters or underlying biological structures. Briefly, the noise value at each pixel was computed by interpolating the dot products between random gradient vectors $g_{i,j}$, and the relative position (u, v) within the grid cell overlaid on the original image. For a given pixel $(x, y)$, the relative position within the grid cell is calculated as,

$$u = x - i, v = y - j, \tag{20}$$

where $i = \lfloor x \rfloor$, and $j = \lfloor y \rfloor$. The dot product between the gradient vectors at the grid's corners and the distance vectors to the pixel can be calculated as,

$$s_{d_x, d_y} = g_{i+d_{x,}, j+d_y} \cdot (u - d_{x,}, v - d_{y,}), \tag{21}$$

where $d_x$, $d_y \in \{0,1\}$, and $g_{i+d_{x,},j+d_y}$ represents the gradient vector at the grid point $(i+d_x, j+d_{y,})$. The final noise value $N(x,y)$ is obtained by linearly interpolating these dot products along both *x*- and *y*-axes using a smooth fade function as follows,

$$N(x,y) = \left[s_{0,0} + f(u)\cdot(s_{1,0}-s_{0,0})\right] + f(v) \\ \cdot\left(\left[s_{0,1} + f(u)\cdot(s_{1,1}-s_{0,1})\right] - \left[s_{0,0} + f(u)\cdot(s_{1,0}-s_{0,0})\right]\right). \quad (22)$$

The fade function is defined as $f(\mathrm{t}) = 6t^5 - 15t^4 + 10t^3$, where *t* represents the relative position within the grid cell along either the x- or y-axis ($u$ or $v$). We randomly combined noise generated using grid cells of varying sizes to create a multi-scale texture that reflects the structured noise consisting of different frequencies.

##### 4.4 Motion-blurred PSFs

To enable SPTnet to recognize fast-moving particles captured under a camera with finite exposure time, we incorporated motion-blurred PSFs into the training process. Starting with a fBm trajectory $\{B_H(T_m)\}$, the exposure time (sampling interval) $T_{exp}$ is divided into *N* intervals of duration $\Delta t$ to generate a finer trajectory $\{B_H(t_n)\}$ with shorter exposure times, where $t_n = \frac{T_m}{N}$. At each time step of $t_n$, the PSF's position can be generated from the scaled $\{B_H(T_m)\}$ based on the self-similarity property of the fBm, which can be expressed as,

$$\{B_H(t_n)\} \cong \{(\frac{1}{N})^H B_H(T_m)\}, \quad (23)$$

where $\frac{1}{N}$ is a scaling factor (set to 0.1), and $\cong$ denotes equal finite dimensional distribution. The blurred PSF is then obtained by summing *N* shorter exposure time PSFs, while maintaining the total intensity the same as that of the PSF with an exposure time of $T_{exp}$.

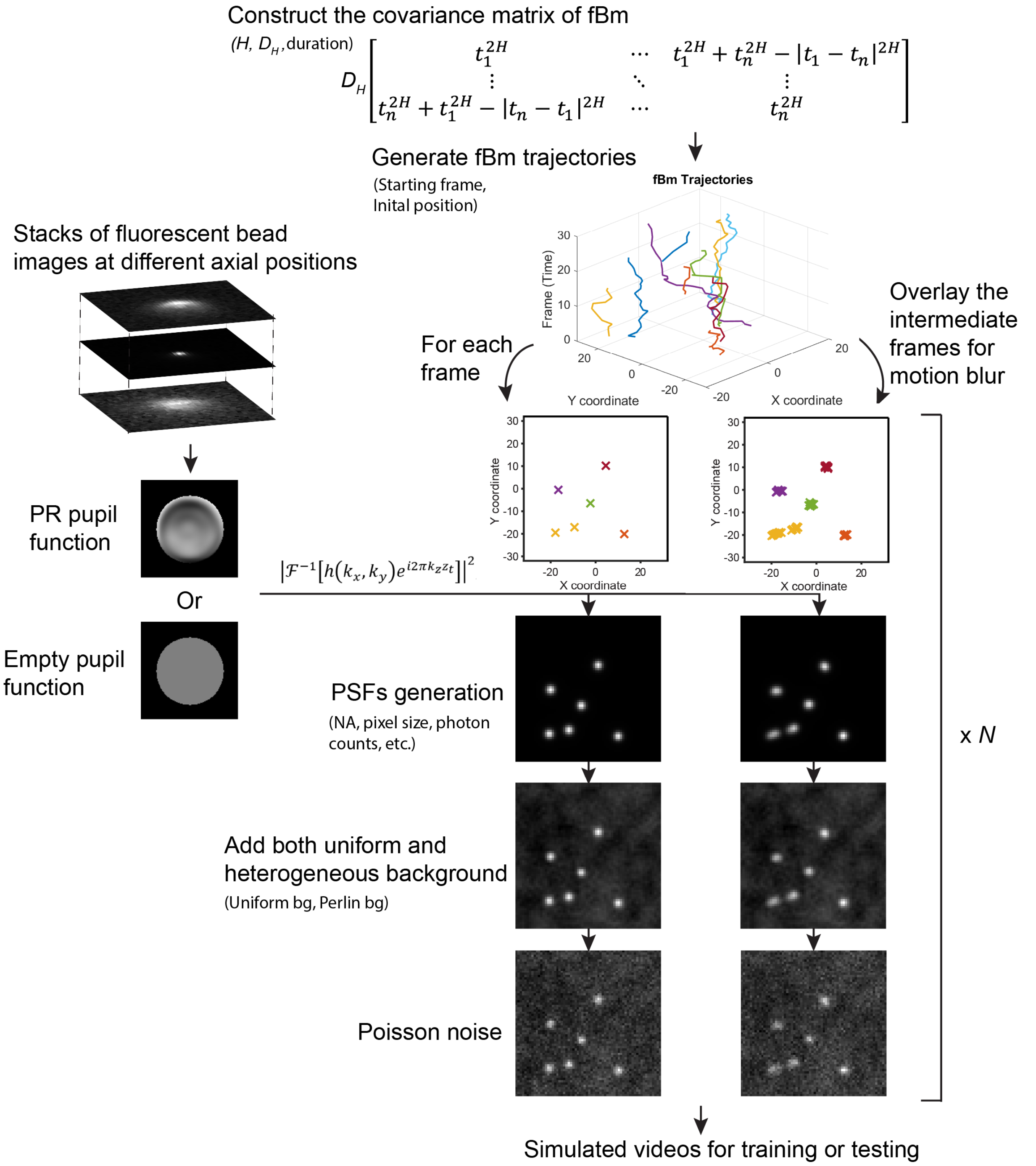

**Fig. SS3: Workflow for generating simulated videos for SPTnet training and testing.** For each particle trajectory in a simulated video, we first construct a covariance matrix based on its Hurst exponent ($H$), generalized diffusion coefficient ($D_H$), and movement duration. Trajectories are then generated via Cholesky decompositions. PSFs at different coordinates are generated through the Fourier transform of the pupil function, which can be obtained using phase retrieval (PR) from the experimental fluorescent bead images at different axial positions, or by using an empty pupil function for systems with minimal aberrations. Motion-blurred PSFs are simulated by overlaying intermediate frames, which are obtained based on the self-similarity of the fBm. Finally, uniform and heterogeneous backgrounds, as well as Poisson noise, are added to each frame to create an *N*-frame simulated video, either with or without motion blur effect.

#### 5. Loss function for simultaneous detection, localization, trajectories construction, and motion parameter estimation.

We constructed a custom loss function to effectively guide the end-to-end learning process for both trajectory prediction and motion parameter estimation. However, given the set-based nature of SPTnet's outputs, direct correspondence between each predicted track query and ground truth labels is not inherently established. To address this, we followed a previously described method[19], employing the Hungarian algorithm to find the optimal one-to-one assignment between predicted track queries and ground truth labels. We minimize the total loss that quantifies the dissimilarity between all Hungarian-matched track pairs. This loss function comprises four different components, as detailed below,

(1). The classification loss ($\mathcal{L}_{cls}$) measures how well the model detects the presence of particles over time. In each simulated video, particles have random starting times and durations. We consider only the particles that last more than two frames as valid tracking targets, assigning a binary class label of 1 for each frame in which they are present. All other cases are labeled as 0 to represent the background. We compute the binary cross-entropy (BCE) loss between the predicted class probabilities $\hat{y}_{cls}$ and the ground truth classes $y_{cls}$,

$$\mathcal{L}_{cls}^{(i,j)} = -\frac{1}{T}\sum_{t=1}^{T}\left[y_{cls,t}^{(j)}\log\hat{y}_{cls,t}^{(i)} + \left(1-y_{cls,t}^{(j)}\right)\log\left(1-\hat{y}_{cls,t}^{(i)}\right)\right], \tag{24}$$

where *T* is the total number of input video frames. $y_{cls,t}^{(j)}$ is the ground truth label for the trajectory *j* at frame *t*, $\hat{y}_{cls,t}^{(i)}$ is the predicted probability that track query *i* represents a particle present at frame *t*.

(2). The Localization loss ($\mathcal{L}_{loc}$) assess the overall localization error of all the particles in each frame. We calculated the pairwise Euclidean distance between predicted position ($\hat{x}_t, \hat{y}_t$) and ground truth position ($x_t, y_t$), masking out any frames where the particle is absent,

$$\mathcal{L}_{loc}^{(i,j)} = \frac{1}{d^{(j)}}\sum_{t=1}^{T}\delta_t^{(j)}\sqrt{\left(\hat{x}_t^{(i)}-\mathrm{x}_t^{(j)}\right)^2+\left(\hat{y}_t^{(i)}-\mathrm{y}_t^{(j)}\right)^2}, \tag{25}$$

where $\delta_t^{(j)}$ is an indicator function that is 1 if the ground truth particle *j* is present at frame *t* and 0 otherwise, and $d^{(j)} = \sum_t \delta_t^{(j)}$ is the duration of the ground truth trajectory *j.*

(3). The Hurst exponent loss ($\mathcal{L}_H$) is the simple mean absolute error (MAE) between the predicted $\widehat{H}^{(i)}$ and ground truth $H^{(j)}$, weighted by the inverse of the scaled Cramér-Rao Lower Bound (CRLB) for Hurst exponent,

$$\mathcal{L}_H^{(i,j)} = \frac{1}{w_H^{(j)}} \left|\widehat{H}^{(i)} - H^{(j)}\right|, \tag{26}$$

where $w_H^{(j)} = \frac{CRLB_H(H^{(j)}, D^{(j)},\ d^{(j)})}{CRLB_H(H^{(j)}, D^{(j)},\ T)}$, and $CRLB_H(H^{(j)}, D^{(j)},\ d^{(j)})$ represents the CRLB for Hurst exponent of a $d^{(j)}$ steps fBm with Hurst exponent of $H^{(j)}$ and generalized diffusion coefficient of $D^{(j)}$.

(4) The generalized diffusion coefficient loss ($\mathcal{L}_D$) is calculated similarly using the CRLB weighted MAE between the predicted $\widehat{D}^{(i)}$ and ground truth $D^{(j)}$

$$\mathcal{L}_D^{(i,j)} = \frac{1}{w_D^{(j)}} \left|\widehat{D}^{(i)} - D^{(j)}\right|, \tag{27}$$

where $w_D^{(j)} = \frac{CRLB_D(H^{(j)}, D^{(j)},\ d^{(j)})}{CRLB_D(H^{(j)}, D^{(j)},\ T)}$, and $CRLB_D(H^{(j)}, D^{(j)},\ d^{(j)})$ represents the CRLB for generalized diffusion coefficient of a $d^{(j)}$ steps fBm with Hurst exponent of $H^{(j)}$ and generalized diffusion coefficient of $D^{(j)}$. These CRLB weightings ensure that tracks with lower theoretical estimation precision are penalized less in the loss function, and also reduce the influence of their motion status estimations in the Hungarian matching process.

The loss function between each prediction and ground truth label is the weighted sum of all four components,

$$C(i,j) = \lambda_{cls}\mathcal{L}_{cls}^{(i,j)} + \lambda_{loc}\mathcal{L}_{loc}^{(i,j)} + \lambda_H\mathcal{L}_H^{(i,j)} + \lambda_D\mathcal{L}_D^{(i,j)}, \tag{28}$$

where the weighting factors $\lambda_{cls}, \lambda_{loc}, \lambda_H$, and $\lambda_D$ were chosen empirically as 1, 2, 0.5, and 0.5, respectively, to balance the scale of each loss component. Each ground truth label is assigned to its optimal matching prediction by the Hungarian algorithm. For the matched pairs, we computed the total loss by summing the loss components for each assigned pair:

$$\mathcal{L}_{match} = \frac{1}{N_{tracks}} \sum_{j=1}^{N_{tracks}} C(\sigma(j), j), \tag{29}$$

where $\sigma(j)$ is the index of the predicted trajectory assigned to ground truth trajectory *j*, and $N_t$ is the total number of ground truth tracks in the video. For all unmatched track queries *i* over T frames, we compute BCE loss,

$$\mathcal{L}_{unmatch} = \frac{1}{(N_q - N_t) \cdot T} \sum_{i=1}^{N_q - N_t} \sum_{t=1}^{T} \log(1 - \hat{y}_{cls,t}^{(i)}), \tag{30}$$

where $N_q$ represents the total number of track queries in SPTnet, and the final cost used for backpropagation is given by,

$$\mathcal{L} = \mathcal{L}_{match} + \mathcal{L}_{unmatch.} \tag{31}$$

#### 6. Conventional algorithms used for comparison with SPTnet in motion parameters extraction

To extract the Hurst exponent *H* and generalized diffusion coefficient $D_H$ from fBm data, several methods are available. In the comparison to SPTnet, we tested some commonly used methods below. While there are other machine learning and deep learning approaches[20-22], they are all based on the analysis of given trajectories and were not specifically designed for optimal performance under the testing conditions of SPTnet, such as short trajectory durations (<30 frames), consideration of localization imprecision, and varying generalized diffusion coefficients. Therefore, for a fair comparison, we focus on comparing SPTnet with conventional estimators.

##### 6.1. Rescaled range analysis (R/S analysis)

Rescaled Range (R/S) analysis was first introduced in 1951 by H. Hurst[23]. It quantifies the variability of a time series by examining how the range of cumulative deviations rescaled by the standard deviation changes with the length of the time series. Briefly, given a *N* steps fBm sample path $\{x_n,\ \mathrm{n} = 1, \dots, N\}$ with a fixed sampling interval, the cumulative deviation can be calculated by,

$$z_t = \sum_{k=1}^{t} y_k \, , t = 1, \dots, N \, , \tag{32}$$

where $y_k = x_n(k) - \bar{x}_n$, is a mean-adjusted time series. and the range R(*N*) is,

$$R(N) = \max_{1 \le t \le N} z_t - \min_{1 \le t \le N} z_t. \tag{33}$$

The Hurst exponent can then be estimated by the linear regression of the logarithm of the following empirical relation,

$$\mathrm{E}\left[\frac{R(N)}{S(N)}\right] = cN^H, \tag{34}$$

where *S(N)* is the standard deviation of the original fBm time series of different lengths, and *c* is a constant. In this work, we used a Matlab (MathWorks, Natick, MA) implementation[24] to estimate the Hurst exponent via R/S analysis.

##### 6.2. Wavelet version of second order derivative (wDSOD) method

The wDSOD method is a wavelet-based adaptation[25] of the discrete second-order derivative (DSOD) method proposed by Istas and Lang[26]. They share the same principle by leveraging the scaling properties of quadratic variations based on second-order increments to effectively estimate Hurst exponent *H*. The key difference lies in their operational domains: the DSOD method operates directly on the raw time series data, while the wDSOD processes the coefficients from the wavelet transform of the original data.

Here, we take the DSOD method as an example to illustrate the estimation process. For *N* steps time series $\{x_n,\ \mathrm{n} = 1, \ldots, N\}$ sampled from the fBm process, the second-order increments are,

$$\Delta_\tau^{(2)} x_n = x_{n+2\tau} - 2x_{n+\tau} + x_n, \tag{35}$$

where $\tau$ is the time lag. The quadratic variation for the second-order increments for each $\tau$ can be calculated as,

$$V_{(2)}(\tau) = \sum_{n=0}^{N_\tau} \left(\Delta_\tau^{(2)} X_n\right)^2, \tag{36}$$

where $N_\tau$ is the total number of increments for different time lag $\tau$. Hurst exponent can be estimated through a log-log regression on the following relation with *k* as a constant,

$$\mathrm{E}\left[V_{(2)}(\tau)\right] = k_{(2)} \tau^{2H}. \tag{37}$$

The built-in function (‘wfbmesti’) from the Matlab wavelet analysis toolbox was used to conduct wDSOD estimation.

##### 6.3. MSD analysis

The mean square displacement (MSD) analysis is a simple and widely used approach to extract the diffusion coefficient and the undergoing types of diffusion (anomalous exponent, $\alpha = 2H$). For an equally sampled *N* steps fBm trajectory $\{x_n,\ \mathrm{n} = 1, \ldots, N\}$, the time-averaged mean square displacement (TAMSD) is calculated as,

$$\mathrm{MSD}(\tau) = \frac{1}{N-\tau}\sum_{n=1}^{N-\tau}(x_{n+\tau} - x_n)^2, \quad (38)$$

where $\tau$ is the time lag, and time lags are the set of all possible steps, and $x_n$ is the position of the fBm at step *n.* Both the $H$ and $D_H$ can be estimated through power-law fitting of the following relationship,

$$\mathrm{MSD}(\tau) = 2D_H\tau^{2H}, \quad (39)$$

It is worth noting that fitting the MSD plot performs optimally under prerequisite conditions: the data follows a normal distribution, and each data point should be weighted by the inverse of its variance[27]. However, due to the difficulty of accurately determining the variance for each data point in practice, we proceeded without applying weights. Moreover, the optimal number of MSD data points used for fitting also significantly influences the results and depends on several factors, including localization error, the total length of the trajectory[27], and Hurst exponent[28]. Consequently, it is difficult to fit each dataset using its optimal number of MSD points.

In this study, instead of using all available MSD points or the first 25%, which is the default setting in a widely used Matlab toolbox "@msdanalyzer"[29], we empirically determined the optimal number of MSD points for fitting through simulations. The optimal number of MSD fitting points for 30-frame datasets under a typical SNR condition (with lateral localization precision $\sqrt{CRLB} \approx 20$ nm) is set to be 5, which is based on the evaluation of the estimation RMSE for both $H$ and $D_H$ under different motion types. We found that the lowest error, considering all motion types, is between 3 and 5, and the absolute differences among these choices are insignificant. To account for the additional localization uncertainty potentially introduced by the heterogeneous background in real data, we chose to use the first 5 data points to conduct the fitting (**Fig. SS4**).

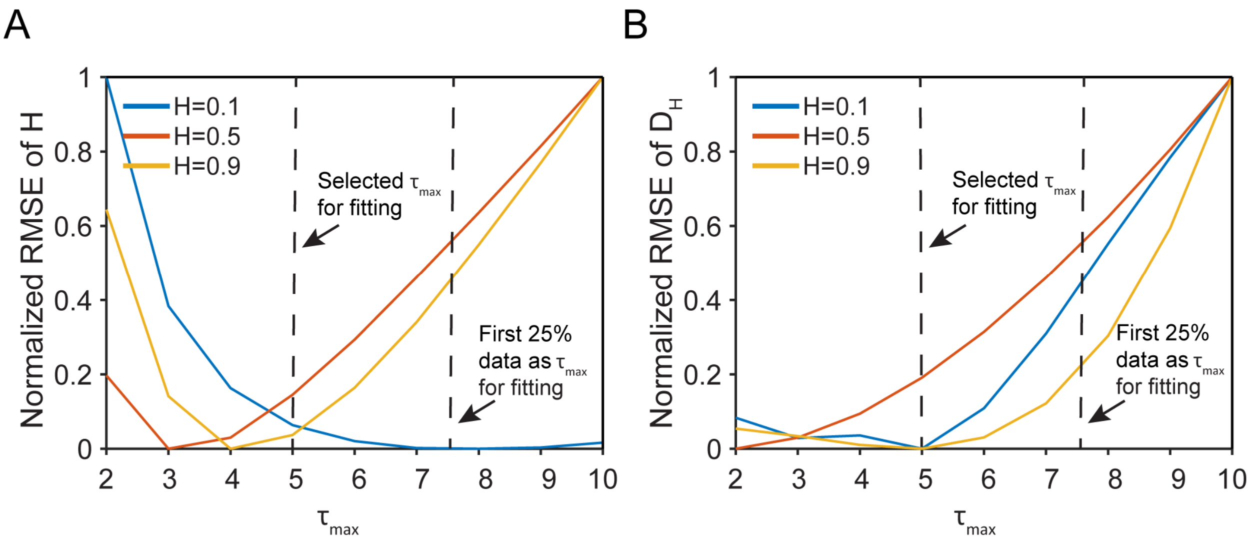

**Fig. SS4: Optimization of the maximum number of time lags ($\tau_{max}$) used for MSD plot fitting**. Simulated fBm trajectories of three motion types, each with 30-frame videos and Poisson noise, were generated using uniformly distributed generalized diffusion coefficients (ranging from 0 to 0.5 pixel$^2$/frame$^{2H}$) and three distinct Hurst exponents. Normalized root mean square errors (RMSE) were calculated for power-law fitting of the MSD plot with varying numbers of $\tau_{max}$. Each RMSE value was obtained from the estimations of 5,000 simulated videos. The dotted line at $\tau_{max}$ = 5 indicates the selected value based on the overall consideration of the RMSE for both Hurst exponent (A) and generalized diffusion coefficient (B) across different motion types. The RMSE obtained by fitting the first 25% of time lags is indicated by the dotted line on the right.

#### 7. Comparison with TrackMate

To the best of our knowledge, no existing end-to-end SPT algorithm provides both trajectories and estimated motion parameters directly from camera images, enabling fair comparisons with SPTnet. TrackMate[30], a widely used toolbox integrates various algorithms for each step of the SPT analysis, allowing comparison of trajectory formed through different processing pipelines. Among TrackMate's built-in detectors, we focused on two commonly used options, excluding simple thresholding and computationally simplified alternatives. The Laplacian of Gaussian (LoG) detector identifies blob-like structures using second-order derivatives with Gaussian smoothing, making it robust to noise but may overlook smaller features. In contrast, the Hessian detector is effective for detecting corners and ridges, though it is more sensitive to noise. The trajectories were formed using the built-in LAP tracker[31], which considers more global information compared to Kalman filter based local optimization approaches. After sub-pixel localization, TrackMate generated trajectories were analyzed through the MSD method to extract Hurst exponent and generalized diffusion coefficient.

A key challenge in this comparison is that, unlike SPTnet, TrackMate requires parameter tuning, and its performance is highly sensitive to these settings. In practice, optimizing parameters for each test video can be difficult. For our comparisons, we manually optimized TrackMate's parameters using the first five examples from each condition (Supplementary Table 4). We also fixed the photon counts and uniform background intensity across all test videos to ensure consistency during parameter optimization. Each test video contained 1 to 5 randomly moving particles present throughout the entire video duration.

##### 7.1 Metric used in comparison

We first employed a linear assignment algorithm to match the detected particle positions to the ground truth positions, using a pairwise Euclidean distances threshold of 3 pixels (pixel size: 157 nm for the TIRF system). Matched predictions were defined as correctly detected molecules, while unmatched ground truth labels were classified as missed detections, and unmatched predictions were classified as spurious detections. Detection accuracy was then quantified using the Jaccard index (JI),

$$\mathrm{JI} = \frac{TP}{FP + TP + FN}, \tag{40}$$

This metric accounts for correctly detected molecules (true positives, TP), spurious detections (false positives, FP), and missed detections (false negatives, NP).

To evaluate the motion parameter estimations, let $r^{(j)}_{pred,i}$ and $r^{(j)}_{true,i}$ denote the predicted and ground truth coordinates at frame *i* of trajectory *j*, respectively. The root mean square error (RMSE) for the Hurst exponent and the generalized diffusion coefficient is calculated only for trajectories where the predicted coordinates remain within a Euclidean distance of $\varepsilon$ from the ground truth at every frame,

$$\begin{gathered} \mathrm{S} = \{ j \mid \|r_i^{pred,j} - r_i^{true,j}\| \le \varepsilon, \forall i \in \{1,2, \dots T_j\} \}, \\ RMSE_H = \sqrt{\frac{1}{N}\sum_{j\in S}(H^{(j)}_{pred} - H^{(j)}_{true})^2}, \\ RMSE_D = \sqrt{\frac{1}{N}\sum_{j\in S}(D^{(j)}_{pred} - D^{(j)}_{true})^2}, \end{gathered} \tag{41}$$

$T_j$ is the total number of frames for trajectory *j*, and $\varepsilon$ is the distance threshold, which is the same as the detection distance threshold. Let *S* be the set of trajectories that satisfy this distance threshold condition, and *N* represents the total number of trajectories in *S*. $H^{(j)}_{pred}$ and $H^{(j)}_{true}$ denote the predicted and ground truth Hurst exponents for trajectory $j$, while $D^{(j)}_{pred}$, and $D^{(j)}_{true}$ are the corresponding generalized diffusion coefficients.

#### 8. Workflow and postprocessing for SPTnet

SPTnet requires minimal input handling and postprocessing. For simulated video, the pixel intensities in each frame are normalized based on the minimum and maximum intensity values across all frames. The normalized video can then be directly analyzed by SPTnet, with a processing time of approximately 60 ms for a 30-frame video covering a 10 μm² field of view (computation time estimated on a GTX1070 GPU). The procedure is similar to experimental data. An additional step is to convert the ADU (analog-to-digital unit) count of each pixel to photon counts. This conversion is based on the camera offsets and gain values determined through experimental EMCCD calibration[32].

SPTnet's output predictions are organized according to the order of track queries, where each query uniquely matches with either a moving particle or the background. The presence of a trajectory in a given frame is determined by applying a detection probability threshold to the output binary classification probability. For most signal-to-noise (SNR) conditions, the detection threshold is set to 0.9. However, for noisy experimental data, we increase this threshold to 0.95 to reduce false positives and ensure higher data fidelity under complex experimental conditions. Notably, the distribution of predicted classification probabilities by SPTnet shows two distinct peaks near 0 and 1, with only 0.21% of predictions falling between 0.1 and 0.9. Moreover, we found the choice of the detection probability threshold has minimal influence on the final detection accuracy (**Fig. SS5**).

The generalized diffusion coefficient is a scaling factor that characterizes the rate of diffusion in anomalous motion system. Unlike Brownian motion, where the mean squared displacement (MSD) scales linearly with time, in fractional Brownian motion, the rate at which particles spread depends on the Hurst exponent. SPTnet outputs generalized diffusion coefficients $D_{SPTnet}$ in simulation units of $pixel^2 \cdot frame^{-2H}$. To convert the network outputs to physical units in $\mu m^2 \cdot s^{-2H}$, we use the following relation,

$$D_{phys} = D_{SPTnet} \cdot {S_P}^2 \cdot (t_f)^{-2H}, \quad (42)$$

where, $S_P$ is the pixel size in micrometers per pixel $(\mu m \cdot pixel^{-1})$, and $t_f$ is the frame time in seconds per frame $(s \cdot frame^{-1})$, $H$ is the Hurst exponent.

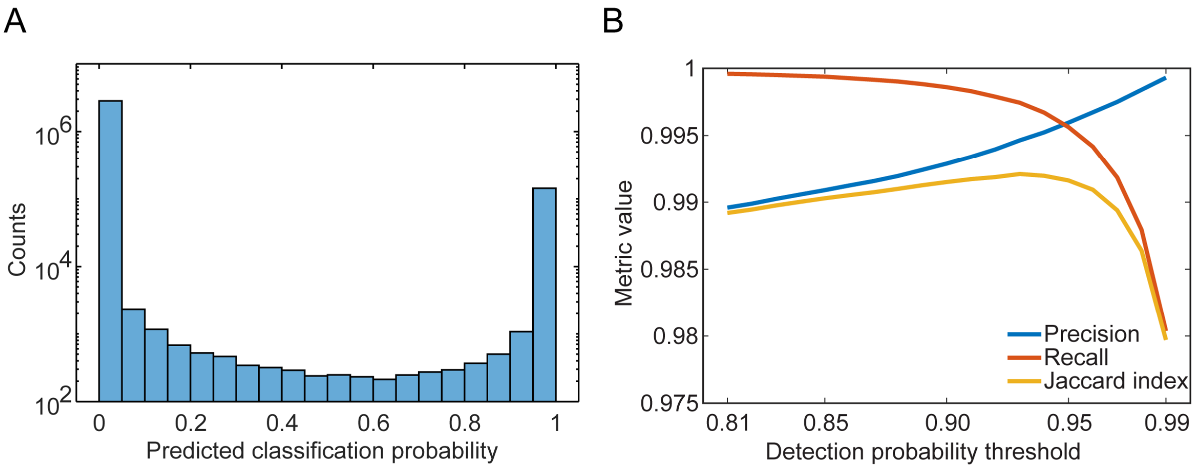

**Fig. SS5: Distribution of SPTnet predicted classification probabilities and the impact of threshold settings on detection accuracy in post-processing.** (**A**) The histogram with a log-scaled y-axis, shows the distribution of predicted classification probabilities from 5,000 videos analyzed by SPTnet. Out of 3,000,000 predictions, 6,393 fall between 0.1 and 0.9. (**B**) Increasing the probability threshold settings in post-processing results in lower recall (TP/(TP+FN)) and higher precision (TP/(TP+FP)). The overall detection accuracy, evaluated using the Jaccard index (TP/(TP+FN+FP)), varies by approximately 1% across the range of tested thresholds.

#### 9. Analysis of sptPALM data in live COS-7 cells

##### 9.1 SPTnet processing of sptPALM data

The current model is optimized for analyzing smaller data blocks of 64 × 64 pixels over 30 frames. To process the larger dataset of 256 × 256 pixels with 1,000 frames, the video was divided into smaller blocks by shifting 32 pixels along both the x and y dimensions to cover the entire spatial area. The temporal dimension was also segmented into 30-frame segments. After SPTnet processed each sub-block, the results were reassembled. We used a detection threshold of 0.95 for all the data, and trajectories with displacements larger than 5 pixels between consecutive frames were considered false linkages and removed. Additionally, repeated detections from overlapping regions were eliminated using a proximity-based threshold. Finally, only tracks with durations exceeding 10 frames were displayed to ensure reliable analysis.

##### 9.2 Generation of the ER tubule centerline

To evaluate the molecules' movement relative to the ER tubule direction, we used a custom workflow in Matlab to find the centerline skeleton to represent the structure of the ER tubules. First, the grayscale image was binarized using adaptive thresholding ("imbinarize" function) with a sensitivity of 0.9, adjusting to local contrast to capture fine details of the ER tubule and minimize background noise (**Fig. SS6, middle**). Next, we applied skeletonization using the "bwskel" function, which reduced the binary ER structures to single-pixel-wide centerlines. A minimum branch length of 5 was set to exclude short, spurious branches, preserving only the main ER pathways. The skeletonized image was then processed with the "bwareaopen" function, applying a threshold of 20 pixels to remove small artifacts and isolated noise, ensuring that only continuous segments of the ER tubule were retained. We then obtained the ER centerline coordinates along the skeleton (**Fig. SS6, right**). These coordinates were fitted with polynomials using the "polyfit" function, which captured the tubule's shape and curvature while smoothing minor variations.

##### 9.3 Jump angle analysis

To quantify deviations of particle trajectories from the endoplasmic reticulum (ER) centerline, we calculated the angle between each trajectory segment and the tangent of the ER centerline at the nearest point. For each trajectory, we identified consecutive points to compute the step vector. We then determined the closest point on the fitted ER centerline and approximated the tangent vector at that location. By normalizing both vectors and calculating their dot product, we obtained the jump angle between the trajectory direction and the ER tubules.

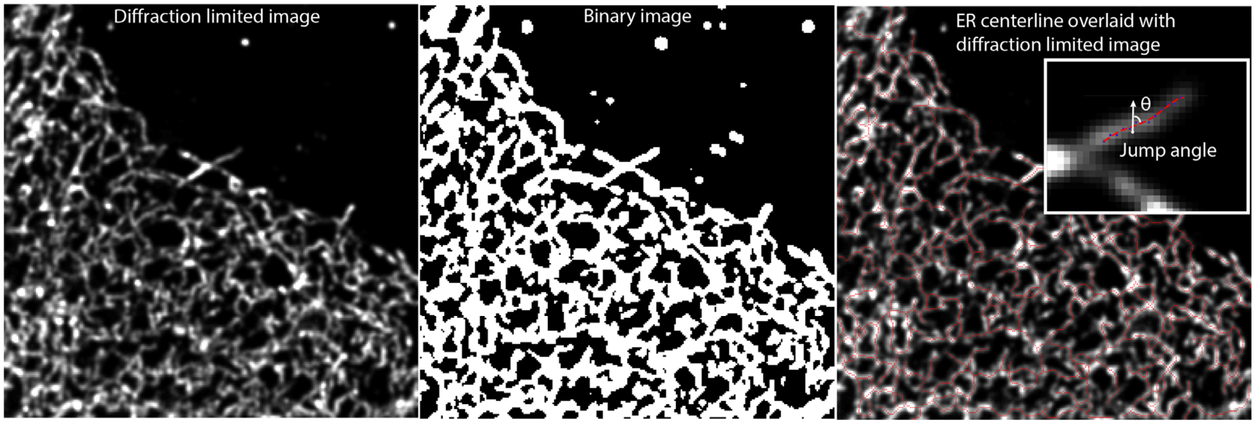

**Fig. SS6: ER centerline generation and jump angle measurement.** The workflow for generating the ER centerline. The inset shows an example measurement of the jump angle (θ) relative to the nearest position on the fitted ER centerline indicated by the red dotted line, at a specific displacement.

#### 10. Apply SPTnet to the 2nd Anomalous Diffusion (AnDi) Challenge.

We evaluated the performance and general usability through the 2nd AnDi challenge[33]. SPTnet was not originally designed to address the specific requirements of the AnDi Challenge. As such, we adjusted the data structure and video frame dimensions to ensure compatibility with our original model. We evaluated SPTnet performance in the two most relevant scenarios: the single-state model (SSM) and the multi-state model (MSM). SPTnet focuses on analyzing apparent motion behavior in videos with minimal constraints and assumptions. While it does not inherently account for dimerization or quenched-trap models, these events, which influence observed motion behavior, can still be inferred from SPTnet outputs. For example, dimerization events can be detected when molecules move closer together and exhibit a change in the diffusion coefficient or anomalous exponent, while environment-induced trapping can be identified by analyzing the average local Hurst exponents.

**10.1 Simulation video generation**

We trained SPTnet using 100,000 simulated videos generated with the AnDi challenge Python package 2.1.5. Tracks are generated by the "models_phenom" class provided in the package. For the MSM model, the probability of switching motion types is defined the same as in the AnDi challenge through the transition matrix M. $M_{i,j}$ represents the probability of switching from state *i* to state *j* at each step. To ensure an unbiased evaluation of SPTnet's performance across various parameter combinations, we tested tracks generated with uniform distributions for both the anomalous exponent and the generalized diffusion coefficient (**Table SS1**), rather than a Gaussian distribution used in the AnDi challenge**.** In this simulation, we directly use parameters to infer the motion states, instead of manually defining the motion states as in the AnDi challenge (e.g., classifying $\alpha > 1.9$ as directed motion) to calculate the classification error. All video-related parameters, such as photon count, numerical aperture (NA), and pixel size, were set to their default values in the "transform_to_video" function provided by the AnDi challenge package.

**Tabel SS1: Parameters of the training and evaluation dataset**

| Model | Numb. Of states | Distribution of Generalized diffusion coefficient | Distribution of anomalous exponent alpha | Model-specific parameters |
|---|---|---|---|---|
| SSM | 1 | Uniform (0,1) | Uniform (0,2) | – |
| MSM | 2 | Unifrom (0,1) | Uniform (0,2) | $M = \begin{pmatrix} 0.99 & 0.01 \\ 0.01 & 0.99 \end{pmatrix}$ |

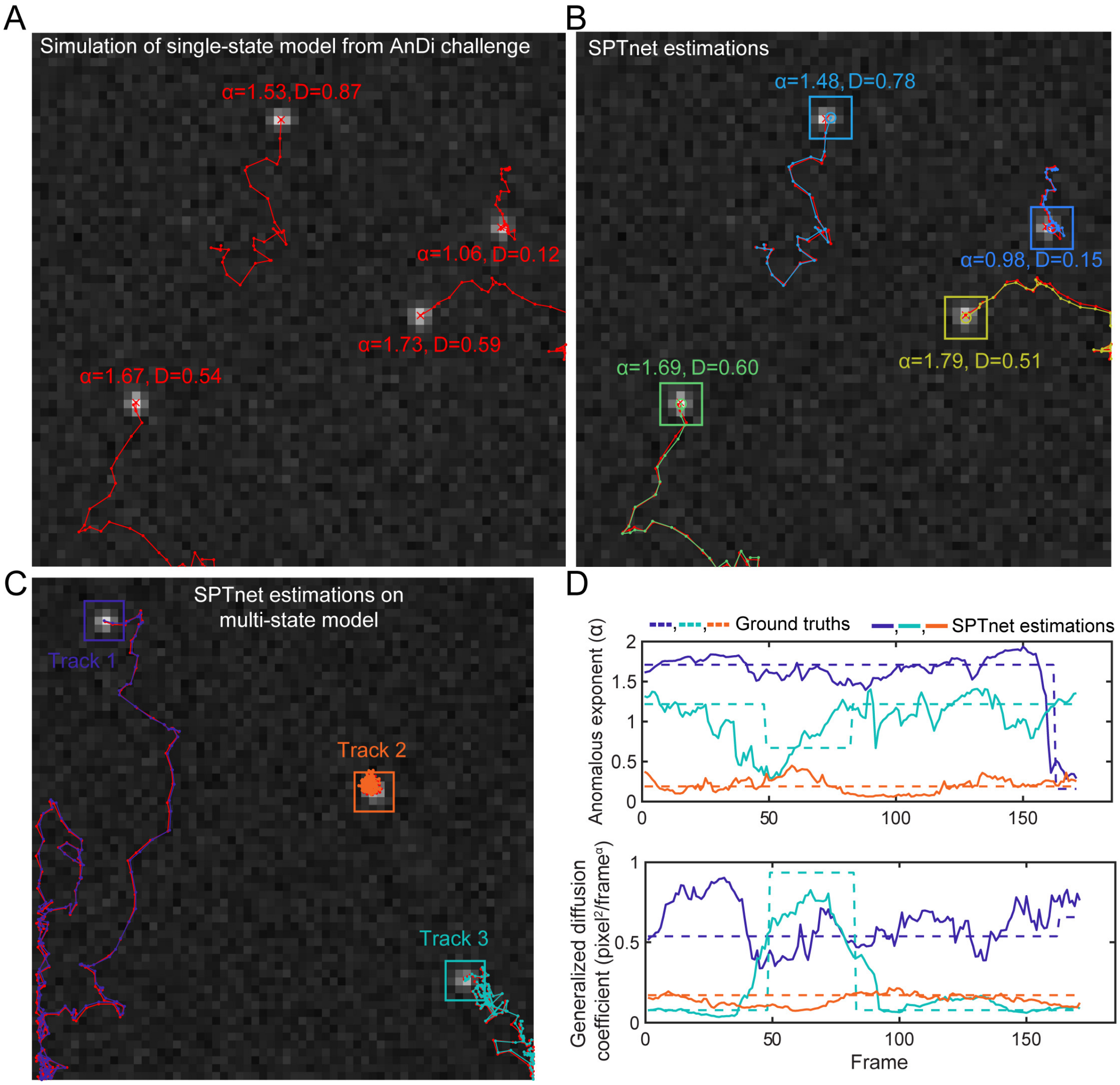

**Fig. SS7: Simulation videos based on the AnDi challenge and corresponding inference results using SPTnet.** (**A**) An example simulation video of the single-state model trajectories, with their corresponding parameters shown in red. (**B**) SPTnet estimation results overlaid with the ground truth trajectory, and the estimated motion parameters displayed alongside detections for each track. (**C**) SPTnet estimated trajectory overlaid with the ground truth of the multi-state model (**D**) The anomalous exponent and generalized diffusion coefficient estimated by SPTnet are plotted against frames for the corresponding tracks shown in (C).

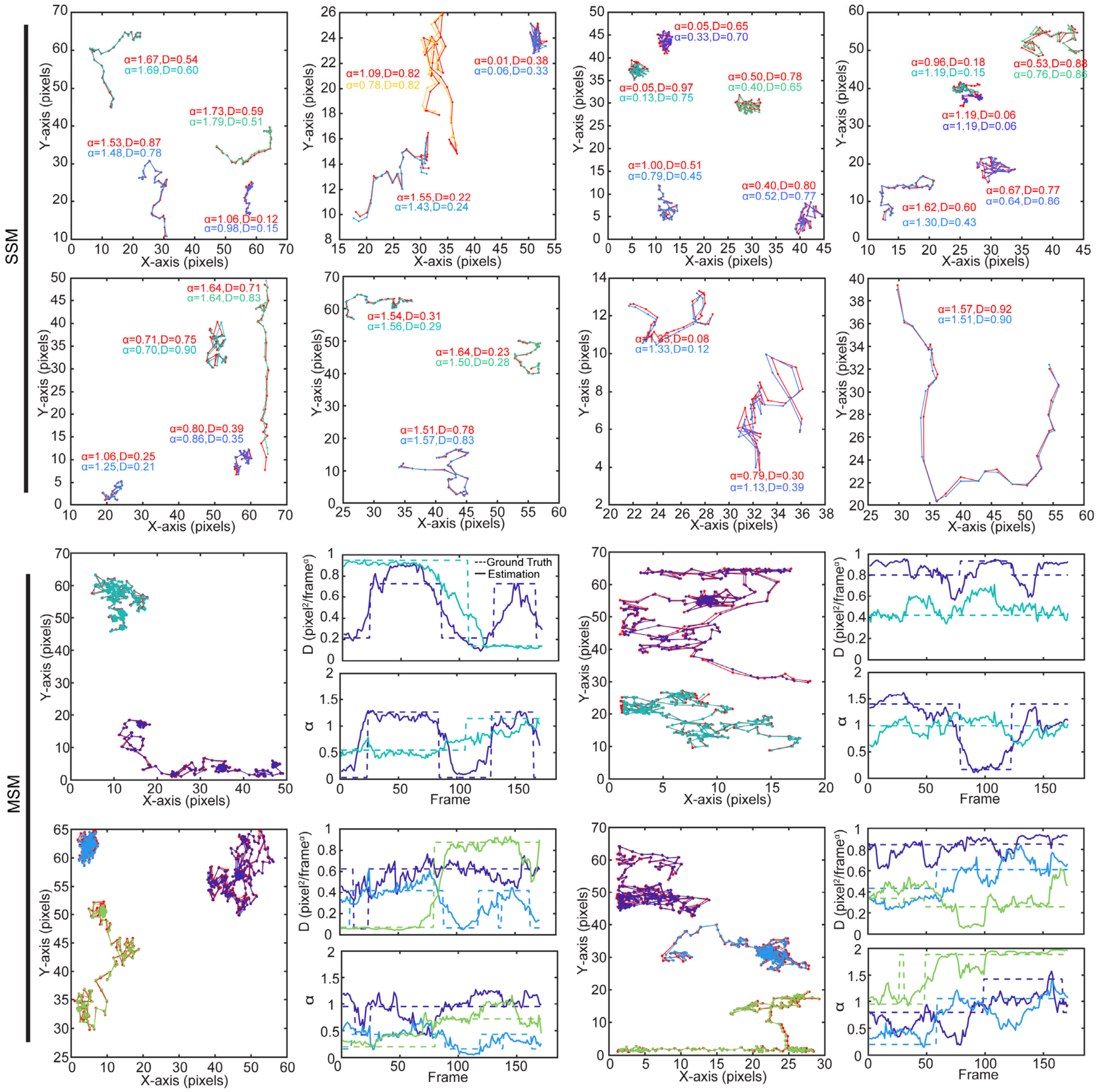

**Fig. SS8: Example results from SPTnet estimations for the single-state model (SSM) and multi-state model (MSM) in the AnDi Challenge.** For the SSM, the ground truth trajectory, anomalous exponent, and generalized diffusion coefficient are shown in red for each track. Below each ground truth, SPTnet's estimated parameters are displayed, and the estimated trajectories are overlaid on the plot. For MSM, SPTnet uses a 30-frame running window for parameter estimation. Changes in motion type across frames are indicated by the dotted line, while SPTnet's estimations are shown as solid lines in corresponding colors for each track.

##### 10.2 Metric used for evaluation

We evaluated the position of each estimation against the ground truth position using the root mean square error (RMSE). Detection accuracy was assessed using the Jaccard similarity coefficient (JSC), defined as in Equation (40). For the anomalous exponent and generalized diffusion coefficient, we adopted evaluation metrics similar to those used in the AnDi Challenge: mean absolute error (MAE) (43) and mean square logarithmic error (MSLE) (44), respectively. However, with a minor modification: instead of evaluating paired change points (CP), we assessed every time step for detections with a localization error smaller than 2 pixels. (**Fig SS7 and Fig SS8**)

$$MAE = \frac{1}{T} \sum_{\substack{paired \\ detections}} \left| \alpha_{gt}(t) - \alpha_{SPTnet}(t) \right|, \tag{43}$$

$$MSLE = \frac{1}{T} \sum_{\substack{paired \\ detections}} \left| \alpha_{gt}(t) - \alpha_{SPTnet}(t) \right|, \tag{44}$$

Since SPTnet does not directly output the CP, for MSM tasks, we inferred the long trajectory using a running window approach. The inferred motion parameters were defined as the values at the middle frame of each running window, resulting in a continuous parameter estimation from frame 15 to frame 185 in a 200-frame video. Unlike the metric used for CP detection, we directly compared the value differences between the ground truth and the estimations for each frame. We use the Hungarian algorithm to match and merge the estimation results from the running windows, taking the median value for repeated localizations. SPTnet achieves the lowest MAE and MSLE for motion parameter estimations (**Tabel SS2**) compared to other methods on the leaderboard. It is worth noting that this comparison is limited to the two scenarios compatible with the original SPTnet output structure and does not account for binding interactions or environmental confinement.

**Tabel SS2: SPTnet evaluation metrics summary**

| | RMSE (localization error) | JSC (detection accuracy) | MAE (anomalous exponent) | MSLE (generalized diffusion coefficient) |
|---|---|---|---|---|
| SSM (single-state model) | 0.328 | 0.966 | 0.145 | 0.007 |
| MSM (multiple-state model) | 0.252 | 0.758 | 0.211 | 0.014 |

**Supplementary Video 1: SPTnet tracking results for spatially overlapping moving particles.**

Cropped subregions from six example input videos each show two transiently overlapping particles, tracked across 30 frames by SPTnet. Each particle has a set of randomly assigned Hurst exponents, generalized diffusion coefficients, and photon counts. Estimated coordinates for each particle are marked in red or blue per frame. At the end of the video, ground truth trajectories appear as dotted lines in the corresponding colors on the line chart.

**Supplementary Video 2: Example comparison between SPTnet and TrackMate under high heterogeneous background.**

Ground truth trajectories of each moving particle are shown as red lines, with a cross indicating the position in the current frame. Estimated trajectories are displayed in distinct colors for each track, with circles marking the estimated positions in the current frame. The heterogeneous background intensity level is set to 40, and TrackMate settings are detailed in **Supplementary Table 4**. For the nearest-neighbor tracker, a maximum linking distance of 6 pixels was used. Scale bars: 500 nm.

**Supplementary Video 3: Example of using SPTnet on experimental supported lipid bilayers (SLBs) tracking data.**

SPTnet's analysis of the diffusion of biotinylated DOPE, labeled with streptavidin Alexa Fluor 647 conjugate. SPTnet estimated trajectories are displayed in different colors, with motion parameters shown at the lower right of each detection box per frame. In panel 3 and 4, a neutral density (ND) filter is implemented to further reduce excitation laser power. Only tracks with detections exceeding 10 frames are shown. Red arrows indicate lipids exhibiting confined movement. Scale bars: 500 nm.

**Supplementary Video 4: Motion behavior switching captured by SPTnet using running window.**

SPTnet's analysis of motion behavior switching using a 30-frame running window with 1-frame step size on the 300-frame videos. In each video, two particles frequently and independently change their Hurst exponent ($H$) and generalize diffusion coefficient ($D_H$). SPTnet's estimates of $H$ and $D_H$ for each molecule are shown in the panels to the right, with colors representing the different molecules. At the end, ground truth trajectories appear as yellow dotted lines, and the ground truth motion parameters are indicated by black dotted lines. Scale bars: 1 µm.

**Supplementary Video 5: Raw sptPALM video of Rtn4 in a live COS-7 cell.**

Raw sptPALM data of Rtn4b-HaloTag labeled with PA-JF$_{549}$-HaloTag ligand in a live COS-7 cell. Scale bar: 5 µm.

**Supplementary Video 6: Raw sptPALM video of Sec61β in a live COS-7 cell.**

Raw sptPALM data of Sec61β-HaloTag labeled with PA-JF$_{549}$-HaloTag ligand in a live COS-7 cell. Scale bar: 5 µm.

**Supplementary Video 7: Example sptPALM video of Rtn4 in a live COS-7 cell analyzed by SPTnet.**

An example input video of sptPALM for Rtn4b-HaloTag labeled with PA-JF$_{549}$-HaloTag ligand in a live COS-7 cell. The input video lasts 30 frames with an exposure time of 0.02 s. The data were analyzed using SPTnet, showing identified trajectories larger than 5 frames with estimated H (hurst exponent) and D (generalized diffusion coefficient) showing at the bottom right corner of each detection box. The detected tracks are overlaid onto a widefield view of the Rtn4 signal.

**Supplementary Video 8: Example sptPALM video of Sec61β in a live COS7 cell analyzed by SPTnet.**

Similar to Supplementary Video 7, this video shows sptPALM of Sec61β-HaloTag. SPTnet identified trajectories show back-and-forth movement along the ER tubules, which is different from the movement pattern of Rtn4.
